# A biofilm-derived peptide as an underwater adhesive

**DOI:** 10.64898/2026.08.05.742774

**Authors:** Xin Huang, Ewelina Liszczyk, Ramesh Prasad, Megan E. Mitchell, Zhijie Wang, Yuchu Liu, Sarvagya Saluja, Mingyu Wang, Rebekah A. Jackson, Peter Dahl, Roberto Andresen Eguiluz, Nikhil Malvankar, Elsa C. Y. Yan, Rich Olson, Huan-Xiang Zhou, Jing Yan

## Abstract

Wet adhesives that perform under water have broad applications in industrial and biomedical settings. To date, molecular designs for underwater adhesives have largely been inspired by marine animals including mussels and barnacles. Here, we propose bacterial biofilms as an alternative source of inspiration for underwater adhesives. Specifically, we demonstrate the application potential of a peptide derived from biofilms formed by the notorious pathogen *Vibrio cholerae*. We characterize the ability of this biofilm-derived peptide to adsorb onto various surfaces and to glue wet surfaces, by using a combination of confocal microscopy, molecular dynamics simulations, atomic force microscopy, lap shear tests, and spectroscopic tools. We further show that the peptide can co-aggregate with microspheres acting as an effective flocculant. Finally, we succeeded in purifying this peptide from *E. coli* in a functional form, setting the stage for large-scale production. Our results open new design possibilities for underwater adhesives inspired by natural biofilms.

**Teaser:** A peptide derived from biofilms adheres to diverse surfaces and shows promising potential as a wet adhesive and flocculant.

## Introduction

Adhesives play important roles in daily life, industrial operations, and biomedical devices (*1*, *2*). They are generally comprised of viscoelastic polymers and are most often applied to clean, dry surfaces. However, many important applications require glues that can be applied and function under wet conditions, such as the repair of submarine vehicles and instruments or the use of tissue adhesives for suture-less wound closure (*2*, *3*). Conventional glues based on synthetic polymers generally perform less well in a wet, complex environment; hence, there is a clear need for new adhesives that can be applied in such conditions. To identify better adhesive materials that function in aqueous environments, extensive studies have been focused on adhesive proteins from various marine animals, particularly mussels, yielding major insights into biological adhesion (*4–8*). The key molecular moiety underlying the adhesive function of mussel foot proteins (Mfps) is L-3,4-dihydroxyphenylalanine (L-DOPA), produced from the amino acid L-tyrosine by the enzyme tyrosinase. The catechol group of DOPA, consisting of a benzene ring with two hydroxyl groups, supports multiple modes of interaction with surfaces, including hydrogen bonding, metal coordination, π-π, amino-π, and cation-π interactions (*9*, *10*). Inspired by the design and adhesive performance of Mfps, purified variations of Mfps as well as synthetic polymers with integrated DOPA units have been used for underwater adhesive applications (*11–17*). However, due to the inherent chemical reactivity of DOPA, Mfps are sensitive to environmental conditions, including oxygen and pH, limiting both their performance in challenging complex environments and our fundamental understanding of the adhesive mechanism. Alternative bioinspired adhesion strategies have also been explored based on barnacles (*18*), ticks (*19*), sea anemone (*20*), starfish (*21*), tunicates (*22*), octopus (*23*), among others.

In this work, we turned to an alternative, untapped source of bioinspiration for underwater adhesion: biofilms. Biofilms are surface-associated bacterial communities encased in an extracellular polymeric matrix (*24–27*). While many bacteria rely on this lifestyle for their survival, biofilms are also widely found in infections and biofouling (*28*, *29*). Indeed, bacteria have evolved for billions of years a surface-associated lifestyle and have consequently developed versatile underwater glues. In particular, *Vibrios* are a major genus among marine bacteria and have adapted to form biofilms on various marine surfaces in salty environments (*30–32*). In a prior study of biofilms formed by *Vibrio cholerae*, the causal agent of the pandemic cholera (*32*, *33*), we found that two partially redundant matrix proteins, Bap1 and RbmC, behave as double-sided tape to anchor *V. cholerae* biofilm clusters to host and environmental surfaces (*34*, *35*). Specifically, both proteins bind to the major biofilm matrix component in *V. cholerae* biofilm, <u>V</u>ibrio <u>p</u>oly<u>s</u>accharide (VPS), via a conserved β-propeller domain while containing a diverse set of surface-binding functionalities in their environment-facing domains. Specifically, Bap1 adheres to abiotic surfaces and lipid membranes via a 57-amino acid loop (hitherto called Bap1-57aa) nested in a β-prism domain. In contrast, RbmC possesses several domains targeting O-glycan-containing mucins and complex N-glycans prevalent in host cell-surface proteins (*36*). Together, Bap1 and RbmC play critical roles in enabling *V. cholerae* biofilms to attach to environmental and host cell surfaces (*34*, *35*, *37*), enhancing colonization and potentially pathogenicity of *V. cholerae* (*38–42*).

Bap1-57aa is largely responsible for the ability of *V. cholerae* biofilms to adhere to different substrata (Fig. 1a) (*34*). Importantly, a chemically synthesized peptide with the Bap1-57aa sequence can spontaneously bind to lipid membranes and glass surfaces outside the biofilm context, suggesting that the Bap1-57aa peptide may serve as a generic and readily tunable underwater glue. In this manuscript, we characterize the ability of this biofilm-derived peptide and its variants to adsorb onto various surfaces and evaluate their potential as underwater adhesives, by using a combination of fluorescence microscopy, molecular dynamics simulations, atomic force microscopy, lap shear tests, and spectroscopic tools. We reveal the molecular conformation and motifs that underlie its surface association, and further show that the peptide can efficiently glue microspheres together to form large precipitates, therefore acting as an effective flocculant. Finally, we successfully purified this peptide from *E. coli* in a functional form, setting the stage for large-scale production.

**Figure 1:**
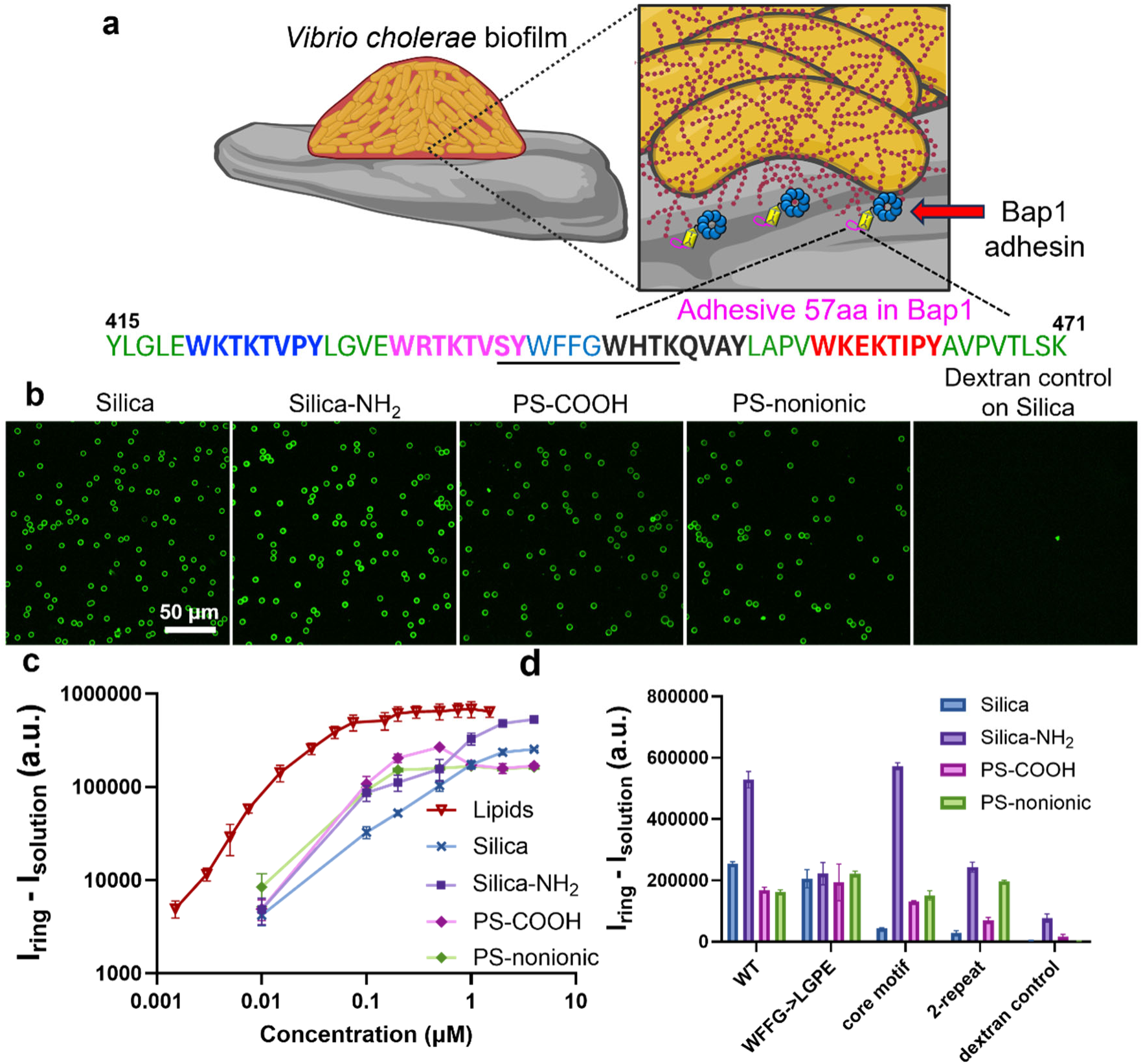
The biofilm-derived Bap1-57aa peptide adsorbs to a wide range of surfaces. (**a**) Schematic of *Vibrio cholerae* biofilms adhering to an abiotic surface. The biofilm-specific adhesion protein Bap1 anchors the biofilm cluster to abiotic surfaces, relying on a 57-amino acid loop (residues Tyr415-Lys471) with the sequence shown. Four pseudo repeats in the Bap1-57aa sequence are shown in blue, magenta, black, and red. The central hydrophobic linker (WFFG) is shown in cyan, and the rest of the residues are shown in green. The core motif of the peptide is underlined. (**b**) From left to right: microscopy images of the FITC-labeled Bap1-57aa peptide spontaneously adsorbing to surfaces of 5 μm silica beads, silica beads with amine groups on the surface (Silica-NH_2_), polystyrene (PS) beads with carboxylate groups on the surface (PS-COOH), and PS beads with nonionic surface coatings (PS-nonionic), respectively. Dextran (4 kDa) conjugated to FITC was used as a negative control. (**c**) Adsorption curves of the Bap1-57aa peptides on 5 μm beads with different chemistries. The excess intensity (I_ring_-I_solution_) was measured for each condition. The lipid adsorption curve was included for comparison. (**d**) Adsorption of various peptide variants on 5 μm beads with different chemistries. Shown are the excess intensity values measured at a peptide concentration of 4 μM (M = mol/L). See Fig. S2 for the definition of variants. Error bars correspond to standard deviation.

## Results

### Quantification of surface adsorption of the Bap1-57aa peptide

Figure 1a presents the sequence of the biofilm-derived peptide. The Bap1-57aa sequence is comprised of four pseudo repeats, with a consensus sequence WbpKpnmY, where b, p, n, and m denote basic (R, K, or H), polar (T, Q, or E), nonpolar (V or I), and mixed (P, A, or S) amino acids, respectively. Basic amino acids occur in two of the eight positions in the pseudo repeats, providing a rich possibility of hydrogen bonds and electrostatic interactions with abiotic surfaces. Aromatic amino acids appear twice in the repeats as well as in the linker, WFFG, between the two inner pseudo repeats.

To systematically quantify the surface adsorption of the Bap1-57aa peptide, we chemically synthesized the peptide with an N-terminal fluorescein isothiocyanate (FITC) label and measured its spontaneous adsorption onto microbeads using fluorescence microscopy (Fig. 1b) (*34*). In brief, 5 μm beads with different compositions and surface properties were incubated with the FITC-labeled peptide and subsequently imaged with confocal microscopy (*43*). The biofilm-derived peptide spontaneously adsorbed onto a range of surfaces, including silica and polystyrene (PS). We also modified the surface chemistry of the beads using either silane chemistry or different PS sources. Bap1-57aa adsorption remained robust across these surface chemistries, highlighting its compatibility with diverse material interfaces and its potential for broad applications.

Subsequently, we generated adsorption curves of the Bap1-57aa peptide on different surfaces by measuring excess fluorescence signals on bead surfaces (I_ring_, “ring” refers to the appearance of fluorescence on confocal images of peptide-bound beads) relative to the solution background (I_solution_) at a series of peptide concentrations (Fig. 1c). The fluorescence signal on the beads is assumed to be proportional to the number of peptide molecules adsorbed on the surface. We generally observed a close-to-linear increase (on a log-log scale) at low peptide concentrations, followed by a plateau at high peptide concentrations, saturating around 0.5-1 μM (M = mol/L). Overall, the peptide exhibits similar levels of adsorption to PS and silica surfaces. Interestingly, modifying the silica surface with amine (silica-NH_2_) increases the adsorption by almost two-fold, possibly due to aliphatic linkages between the amine group and the silica surface, which increase the surface’s propensity for nonspecific adsorption, as can be seen in the dextran-FITC control (Fig. S1). For this reason, we will pay less attention to adsorption results for the silica-NH_2_ surface. For comparison, we have also included an adsorption curve on lipid-coated beads (*44*); binding to lipid bilayers is stronger than to abiotic surfaces, indicating that there are modes of interactions, such as membrane insertion, that are not possible on abiotic solid surfaces.

After establishing the binding assay, we sought to elucidate the surface-association mechanism of Bap1-57aa by designing and testing a series of peptide variants (Fig. 1d). Following our prior work on the peptide-lipid binding (*44*), we studied the following set of mutants: 1) a short peptide with 10 amino acids corresponding to the “core motif” of the peptide (SYWFFGWHTK); 2) a 57aa peptide with the central linker WFFG swapped to LGPE (WFFG→LGPE), preserving the contribution of the four pseudo-repeats while removing the effects of aromatic residues in the central linker; 3) a truncated peptide with only the first two pseudo-repeats (2-repeat; LGLEWKTKTVPYLGVEWRTKTVSY). The full amino-acid sequences of these peptides can be found in Fig. S2.

To simplify data interpretation, we measured and compared surface adsorption signals at a saturating concentration of 4 μM (Fig. 1d). For adsorption on the silica surface, the core motif contributes minimally, which may be related to the hydrophobic nature of the aromatic residues in the core motif and the hydrophilic nature of the silica surface. On the other hand, the pseudo-repeats appear to contribute additively to adsorption through avidity effects. While the 2-repeat sequence has no measurable adhesion (close to the negative controls), the WFFG→LGPE mutant with 4 pseudo-repeats shows an adsorption level similar to the counterpart of the wild-type (WT) Bap1-57aa peptide. Presumably, basic residues in the pseudo-repeats engage in electrostatic interactions with the negative charges on the silica surface. For adsorption on PS beads coated with either carboxylate groups or nonionic, hydrophilic stabilizers, both the core motif and the peripheral pseudo-repeats contribute to adsorption, albeit to different extents.

Overall, our data shows that the biofilm-derived peptide can spontaneously adsorb to a range of surfaces. Mechanistically, the hydrophobic-residue-rich core motif and the peripheral pseudo-repeats can both contribute to surface binding to varying degrees, depending on the surface chemistry.

### Quantification of peptide-surface interactions with a quartz crystal microbalance

The fluorescence-based bead assay allowed us to make quantitative comparisons between different variant peptides; however, it does not give us an absolute measurement of how many molecules are adsorbing onto the surface. Also, it is difficult to test the reversibility of the adsorption process in this assay. To address these limitations, we performed measurements of the WT peptide binding to silica surfaces using a quartz crystal microbalance with dissipation (QCM-D), a sensitive technique for quantitative measurement of surface adsorption and for assessing the reversibility of the adsorption process (*45*, *46*). A SiO_2_ chip was initially rinsed with water and 0.7 % (volume fraction) dimethyl sulfoxide (DMSO), then incubated with 1 μM peptide dissolved in 0.7 % DMSO, and lastly rinsed with 0.7 % DMSO, isopropanol, ethanol, and 10 g/L sodium dodecyl sulfate (SDS) solution (Fig. 2a). After each rinse, values in water were recorded again (Fig. 2a, b). Frequency and dissipation were recorded in the 3^rd^, 5^th^, 7^th^, and 9^th^ harmonics, which had low variability (Fig. S3).

**Figure 2:**
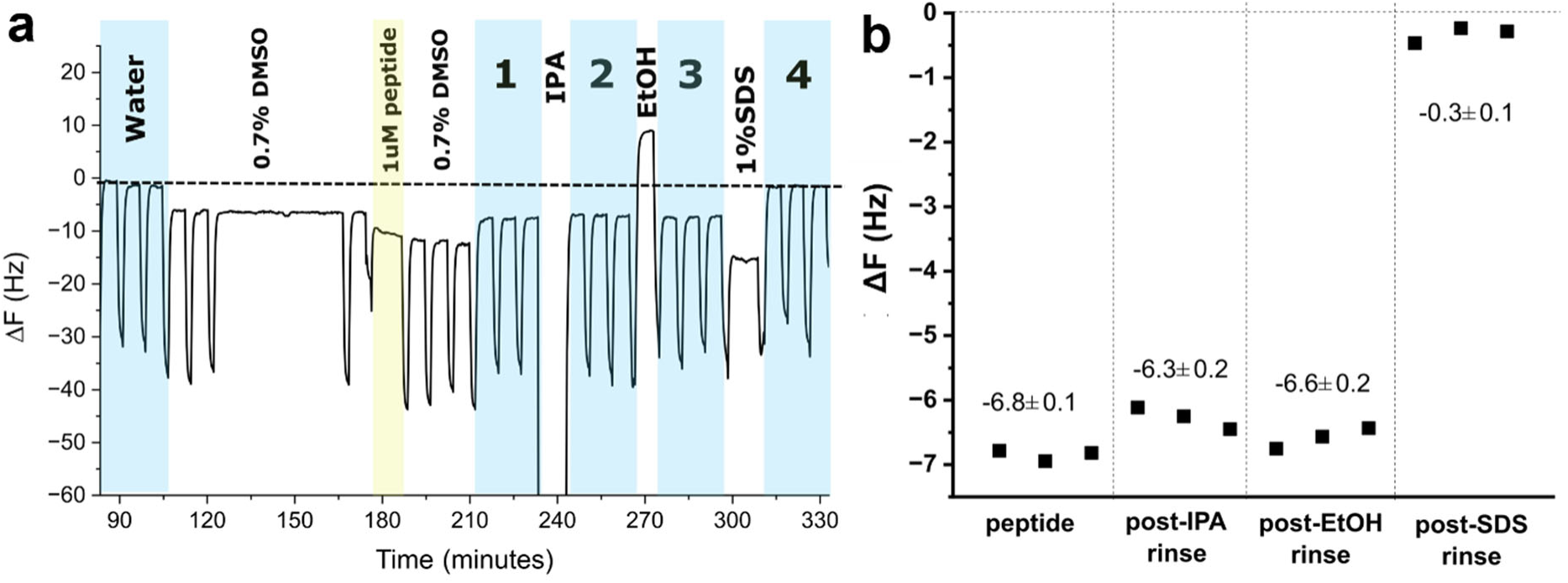
Representative quartz crystal microbalance measurement of Bap1-57aa adsorption on a SiO_2_ chip. (**a**) Changes in 5^th^ harmonic frequency over time relative to the initial water injection. Sharp decreases in frequency are present at the time of injection, likely due to temperature variation upon injection. The yellow shading indicates where the peptide was injected. The dashed line represents the initial water value. Blue shading is used to denote measurements in water. The water measurements labeled 1, 2, 3, and 4 are plotted for clarity in b. (**b**) Difference in frequency for the SiO_2_ chip in water after incubation with the peptide, rinsing with isopropanol (IPA), rinsing with ethanol (EtOH), and rinsing with sodium dodecyl sulfate (SDS). The zero value was set to the average frequency for five measured water injections prior to incubation with peptide. The average of the three measurements is shown above the data points, with the error corresponding to the standard deviation (SD).

The binding of the Bap1-57aa peptide to SiO_2_ is largely irreversible, as the measured frequency when exchanged into pure water after peptide adsorption remained constant at around −6.5 Hz compared to the initial value in water. Even after rinsing with isopropanol or ethanol, the measured frequency of the silica surface with peptide bound did not return to the initial value in water. This stable adsorption is useful for potential underwater applications. After rinsing with SDS, the frequency almost returned to zero, with an average value of (−0.3 ± 0.1) Hz, which is within the sensitivity of the instrument (1 Hz), suggesting that SDS removed most of the peptide from the surface.

We then repeated the experiment four times to assess consistency. The average frequency change in the 5^th^ harmonic of the peptide layer on the surface for each experiment was (−6.7 ± 0.6), (−6.8 ± 0.1), (−4.1 ± 0.1), and (−5.6 ± 1.4) Hz, respectively (average ± standard deviation). The expected sensitivity of the OpenNEXT QCMD instrument with 5 MHz sensors is 1 Hz (or equivalently 17.7 ng/cm^2^). This allows us to estimate the expected signal from the peptide using the Sauerbrey relation (*47*), by assuming a complete, continuous monolayer of protein lying flat on the surface, homogeneous and isotropic. The thickness of this layer is estimated to be the thickness of a polypeptide chain, or 0.8 nm. The estimated density is that of protein, 1.2 g/cm^3^.

The mass per area is thus:

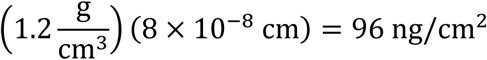

leading to an expected frequency change of −5.5 Hz for a complete monolayer of protein. The average frequency change for our four measurements was (−5.8 ± 1.3) Hz, which agrees with the expected value for a monolayer, corresponding to a 5 % difference between the measurement and the theoretical estimate. We caution, however, that this estimate is based on many assumptions, particularly that the peptide homogeneously covers the silica surface. In practice, strong peptide-peptide interactions can cause aggregation and consequently a heterogeneous film.

### A biofilm adhesion assay confirms the contributions of various motifs in Bap1-57aa

Previously, we showed that the Bap1-57aa peptide is critical for the proper adhesion of *V. cholerae* biofilms to abiotic surfaces, using a biofilm adhesion assay (*34*). Here, we used the same assay to test some of the conclusions from the *in vitro* characterization (Fig. 3a). To do so, we first generated *V. cholerae* mutants containing various *bap1* constructs (*34*). We then grew mature biofilms of *V. cholerae* cells constitutively expressing mNeonGreen on glass surfaces, followed by vigorous washing. Finally, we quantified the biomass on the glass before and after the wash and used the ratio between the two as an indication of adhesion strength. Biofilms possessing a WT 57aa sequence will have a value close to 1, whereas non-adhesive mutants, such as the 57aa deletion mutant (Δ*57aa*), will have a value close to 0. We used increasing concentrations of bovine serum albumin (BSA) as a competitive inhibitor of surface binding.

**Figure 3.**
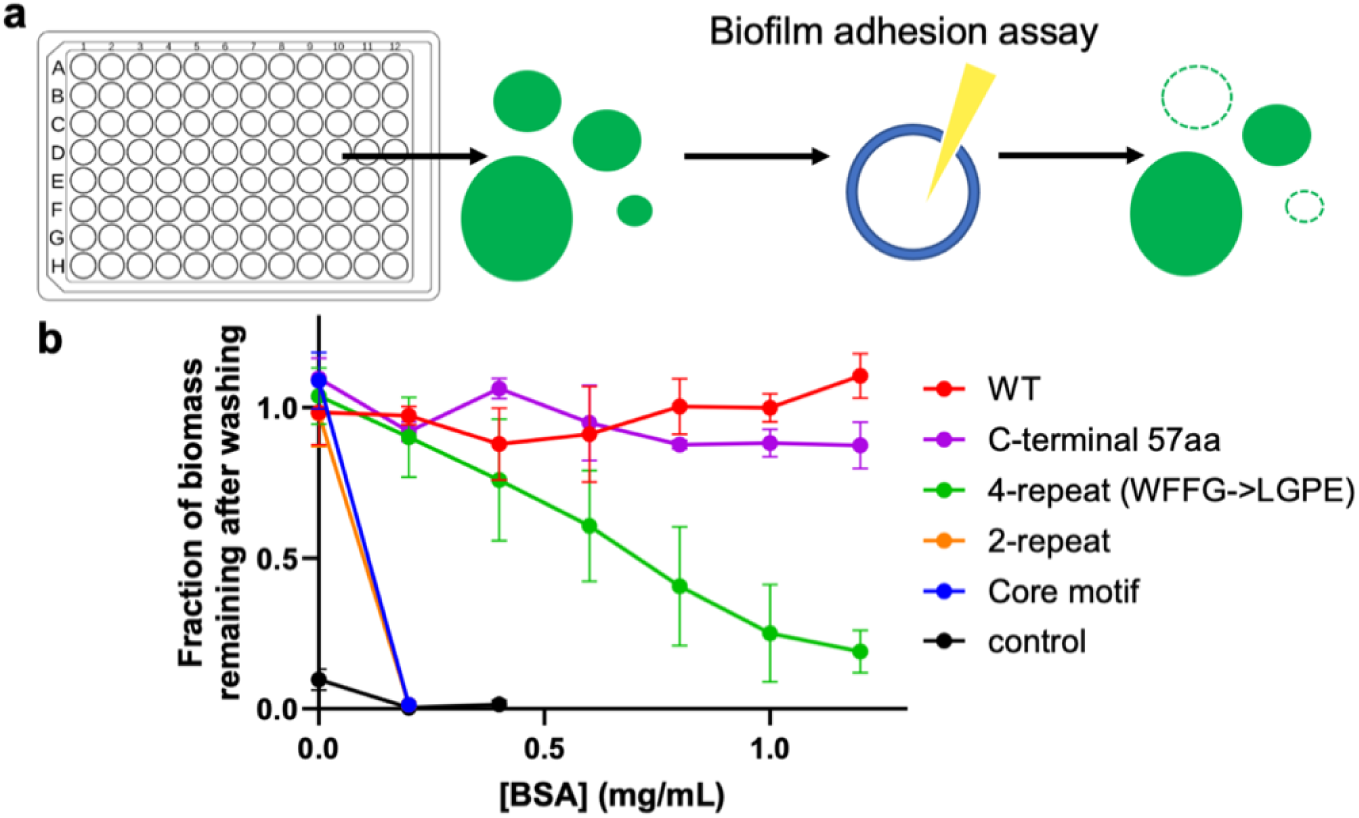
Adhesive function of the Bap1-57aa peptide in the native biofilm context. (**a**) Schematic of the biofilm adhesion assay. Briefly, biofilms from *V. cholerae* cells constitutively expressing mNeonGreen and various *bap1* mutants were grown overnight and imaged with confocal microscopy. Subsequently, the biofilms were vigorously washed and imaged again at the same location. The fraction of biofilm biomass remaining on the substrate reflects the adhesive ability of the biofilms. (**b**) Results from the biofilm adhesion assay for different peptide variants. BSA was used at increasing concentrations during biofilm growth as a non-specific competitor for glass surface adhesion. A Δ*57aa* strain was used as the negative control. All data are depicted as mean ± SD (*n* = 3).

As shown previously, we verified that biofilms with a WT 57aa sequence robustly adhere to the glass surface regardless of the presence of BSA, while the negative control has minimal ability to adhere to glass (Fig. 3b). Next, we tested several 57aa variants with sequences which correspond to the peptides we have chemically synthesized and tested in the bead adsorption assay. Replacing the full 57aa sequence with the 10aa core motif significantly affects the adhesion ability (Fig. 3b, Core motif): the corresponding mutant biofilm lost adhesion as 0.2 mg/mL BSA was added. The mutant biofilm with the 2-repeat peptide is similarly defective in this assay (Fig. 3b, 2-repeat). The 4-repeat variant (WFFG→LGPE), however, showed greatly improved glass adhesion, though still lower than the WT (Fig. 3b, WFFG→LGPE). As an alternative method to test peptide-surface interactions in the biofilm context, we also visualized the spatial localization of Bap1 molecules in biofilms using a 3×FLAG tag combined with a fluorescent Anti-FLAG antibody (*34*, *37*, *48*). Indeed, while WT Bap1 signal concentrates underneath the biofilm, in all defective mutants, the Bap1 surface signals are significantly diminished, consistent with their reduced adhesive abilities (Fig. S4). All these results are consistent with the observations in the *in vitro* bead adsorption assay.

We note a potential difference in the microenvironment of the Bap1-57aa peptide in the biofilm-based and *in vitro* assays: in the former, the 57aa is a loop nested in a well-folded domain, whereas in the latter, the Bap1-57aa peptide lacks end-to-end restraints. To test if this difference affects our conclusions, we constructed a mutant in which the 57aa sequence was repositioned at Bap1’s C-terminus, thus releasing the end-to-end restraints on the peptide. We found that this mutant has an intact ability to bind to the glass surface (Fig. 3b, C-terminal 57aa), suggesting that the 57aa peptide can function independently of its original loop context when adhering to glass – consistent with the *in vitro* results with chemically synthesized peptides that have no such conformational restraints.

### MD simulations reveal the molecular mechanism of surface adsorption

To further understand the molecular mechanism underlying the surface association of the Bap1-57aa peptide, we performed all-atom molecular dynamics (MD) simulations of the peptide on a silica surface. Unlike our previous work on Bap1-57aa-lipid interactions (*44*), we expect that its interaction with the silica surface is largely electrostatic in nature, given the polar and negative charge characteristics of the surface. The Bap1-57aa peptide is enriched in charged residues, with eight positively charged residues (Lys and Arg) and three negatively charged residues (Glu), resulting in a net charge of +5 (Fig. 4a). Indeed, analysis of the contact frequencies of individual residues with the silica surface (with a 3.5 Å distance cutoff) shows that several positively charged (Lys, Arg) and polar (Ser, Trp and Tyr) amino acids frequently form hydrogen bonds with the silica surface (Fig. 4b).

**Figure 4:**
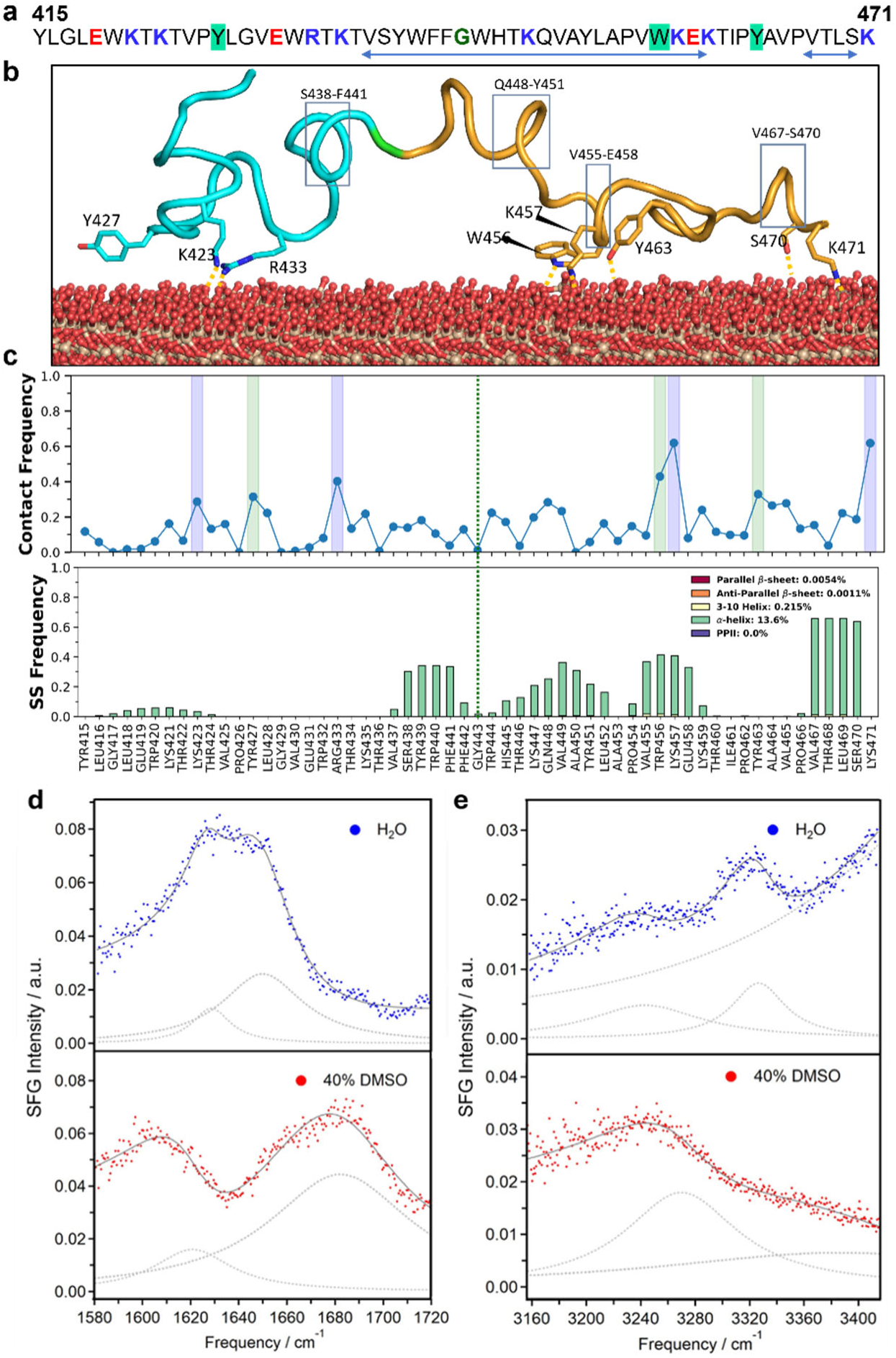
Interaction of the Bap1-57aa peptide with silica surfaces induces helix formation, revealed by molecular dynamics (MD) simulations and sum frequency generation (SFG) spectroscopy. (**a**) Sequence of the Bap1-57aa peptide with aromatic residues involved in surface interactions highlighted by green, positively charged residues in blue, and negatively charged residues in red. The central residue Gly443 divides the Bap1-57aa peptide into N-half (Tyr415-Phe443) and C-half (Trp444-Lys471). The arrow indicates the region of helix formation, most likely located in the C-half of the peptide. (**b**) Representative snapshots from MD simulations of a Bap1-57aa peptide on a silica surface showing contacts with negatively charged surface sites. Interacting residues Tyr427, Lys423, Arg433, Trp456, Lys457, Tyr463, Ser470 and Lys471 are shown in stick representation. The N-half and C-half of the peptide are shown in cyan and orange cartoons, respectively. The silica surface is shown in sphere representation, with O and Si atoms colored red and orange, respectively. Regions with a helical conformation are highlighted with rectangles. (**c**) *Top*: Per-residue contact frequency with the silica surface within a 3.5 Å distance cutoff between peptide and silica heavy atoms. Ten independent simulations were performed. *Bottom*: Frequency of secondary structures (SS) of the Bap1-57aa peptide for each residue from simulations starting from a disordered configuration. Average contact frequencies and SS are calculated from the last 500 ns of each replica, corresponding to a total of 5 µs of simulation time. (**d**) SFG spectra in the amide I region and (**e**) in the NH stretch region of the Bap1-57aa peptide deposited on a glass slide prepared using H_2_O (*top, blue*) and 40 % DMSO (*bottom, red*) as solvents. The spectral fitting is shown in solid grey lines with the fitting component peaks shown in grey dashed lines.

To generate further insights, we divided the Bap1-57aa sequence into N-half (Tyr415-Phe442) and C-half (Trp444-Lys471), with each half containing two pseudo-repeats. Notably, the contact frequency analysis shows that the C-half interacts more strongly with the silica surface than the N-half (Fig. 4b, c top). Although the distribution of positively charged residues is equal in both halves of the peptide, the N-half contains two negatively charged Glu residues (Glu419 and Glu431), whereas the C-half contains only one (Glu458). Moreover, the repulsive effect of Glu458 in the C-terminal half is reduced because it is flanked by two positively charged Lys residues. In contrast, in the N-terminal half, Glu residues are not next to a positively charged residue.

The electrostatic interactions with the surface largely explain the surface adsorption abilities of different peptide variants (Fig. 1d and 3b): the WFFG→LGPE mutant largely retains the charge distribution of the WT Bap1-57aa sequence and hence has a similar or slightly reduced adsorption strength. The 2-repeat has a reduced avidity effect due to the reduced number of positive charges. Finally, the core motif, although showing strong lipid association due to membrane insertion (*44*), performs worse when interacting with various solid surfaces.

In addition to the key residues for surface association, the MD simulations revealed interesting conformational information (Fig. 4b). In our previous work on the Bap1-57aa-membrane interaction (*44*), we showed that in solution, the Bap1-57aa peptide forms a mixture of random coils and β-sheets, which convert into a characteristic β-hairpin structure upon inserting into the hydrophobic core of lipid bilayers when encountering a membrane. In contrast, here we found that, starting from a disordered conformation, several regions of the Bap1-57aa peptide, especially in the C-half, adopt a helical conformation with 35% to 65% probabilities upon interacting with the silica surface (Fig. 4c, *bottom*). The transition from a disordered state to helical formation is driven primarily by the tethering of the residues within (W456 and K457) and flanking (R433, Y463, and K471) the helical regions.

Collectively, these observations demonstrate that binding between the Bap1-57aa peptide and the silica surface is largely regulated by the spatial arrangement of positively and negatively charged residues and hydrogen-bond-forming residues. Our results also highlight how an intrinsically disordered peptide can adopt different conformations when interacting with different surfaces.

### Sum frequency generation spectroscopy confirms the helical conformation of the peptide on silica surfaces

Next, we sought to verify the α-helical conformation of the peptide when adsorbing on negatively charged silica surfaces, as predicted by the MD simulations. Since most spectroscopic techniques, such as circular dichroism, only give information of the molecular conformations in solution, we resorted to vibrational sum frequency generation spectroscopy (SFG), a technique specifically sensitive to vibrational structures of molecules adsorbed at interfaces (*49–52*). In an SFG experiment, an infrared (IR) beam at frequency *ω*_IR_ and a visible beam at frequency *ω*_vis_ overlap temporally and spatially at the sample surface to generate SFG signals at the sum frequency (*ω*_SFG_ = *ω*_vis_+ *ω*_IR_). This optical process depends on the second-order susceptibility, 3(^2^), which is zero in centrosymmetric media, such as bulk liquids. At interfaces, however, centrosymmetry is broken, leading to a non-zero 3(^2^). As a result, SFG is sensitive to molecules adsorbed at interfaces. When applied to study proteins at interfaces, SFG can characterize secondary structures by probing the peptide backbone amide I and/or NH stretching modes (*49*, *50*, *52*). The amide I bands at lower frequencies (1620-1630 cm^−1^) and/or higher frequencies (1680-1690 cm^−1^) are indicative of β-sheet structures, whereas amide I bands in the range of 1640-1660 cm^‒1^ are characteristic of α-helices. Moreover, the NH stretching frequencies are higher for α-helical structures compared to β-sheet structures due to the weaker hydrogen bonds experienced by the NH groups in α-helices (*50*, *52–54*).

To investigate the peptide conformation at the interface, we examined the secondary structures of the Bap1-57aa peptide deposited on a glass substrate. To illustrate the sensitivity of SFG to peptide secondary structures, we collected the amide I and NH spectra (Figs. 4d and 4e) of the sample prepared using pure water and then repeated the experiments with the sample prepared using 40 % DMSO (volume fraction). As the peptide dissolves more readily in DMSO than in pure water (*34*, *44*), we hypothesized that 40 % DMSO will disrupt the surface-peptide interactions and perturb the peptide secondary structures. The SFG spectrum of the sample prepared using pure water (Fig. 4d, top) shows two peaks at (1628.3 ± 0.5) and (1649.9 ± 0.4) cm^−1^, corresponding to β-sheet and α-helical structures, respectively, with the latter showing a higher amplitude in the fitted result (dotted curves in Fig. 4d top; Table S1). These peaks are not due to water OH bending modes, as they remain largely unchanged when H_2_O is replaced by D_2_O (Fig. S5). The SFG spectrum of the sample prepared using 40 % DMSO (Fig. 4d, bottom) shows significantly different amide I bands at (1620.8 ± 1.2) and (1682.2 ± 2.0) cm^−1^, which are highly characteristic of β-sheets (Table S2). Figure 4e shows the NH spectra for the sample prepared using pure water (Fig. 4e, top), which exhibits peaks at (3246.5 ± 4.0) and (3328.0 ± 1.9) cm^−1^, attributed to β-sheet and α-helix structures, respectively (Table S1). Conversely, the sample prepared using 40 % DMSO (Fig. 4e, bottom) shows a pronounced peak at (3269.8 ± 2.9) cm^−1^, attributed to β-sheet structures (Table S2). These experiments clearly show that SFG is sensitive to the secondary structure of the Bap1-57aa peptide adsorbed on the glass surface. The results also reveal that the Bap1-57aa peptide, when first dissolved in water, adopts α-helical conformations on the glass surface with some contribution of β-sheets (Fig. 4d-e, Table S1). Because depositing proteins on solid substrates often induces protein aggregation yielding β-sheet-rich structures (*55*), our observation of Bap1-57aa in α-helical structures is consistent with the MD simulations (Fig. 4c) highlighting the α-helical conformation near the glass substrate.

### Atomic force microscopy and lap shear tests of the adhesive ability of the biofilm-derived peptide

Our characterization so far focused on the adsorption and conformation of the Bap1-57aa peptide. To evaluate its potential as an underwater adhesive, we performed a series of experiments using atomic force microscopy (AFM) to quantify the underwater adhesion strength of the biofilm-derived peptide (Fig. 5a). We have previously used this AFM-based method to measure adhesion of various mutant *V. cholerae* cells to determine the proteins involved in adhesion (*40*). Briefly, silica beads were first attached to tipless AFM cantilevers and subsequently coated with peptide by incubating with 4 mM peptide in water for > 20 minutes. Next, the bead with adsorbed peptides was allowed to approach a pristine mica surface in water, and forces were measured. Upon retraction of the probe from the surface, the force-distance curves demonstrated a substantial adhesive force corresponding to the adhesion conferred by the Bap1-57aa (Fig. 5a, Fig. S6). From the force-distance curves, we extracted the maximum adhesive force or pull-off force. By repeating this measurement multiple times at multiple locations, we generated the distribution of pull-off forces for each peptide variant (Fig. 5b). We observed that the WT peptide shows the strongest pull-off force (11.5 nN), whereas both the core motif and the WFFG→LGPE variant show a more than 3-fold reduction in pull-off force. This observation is largely consistent with the adsorption data.

**Figure 5:**
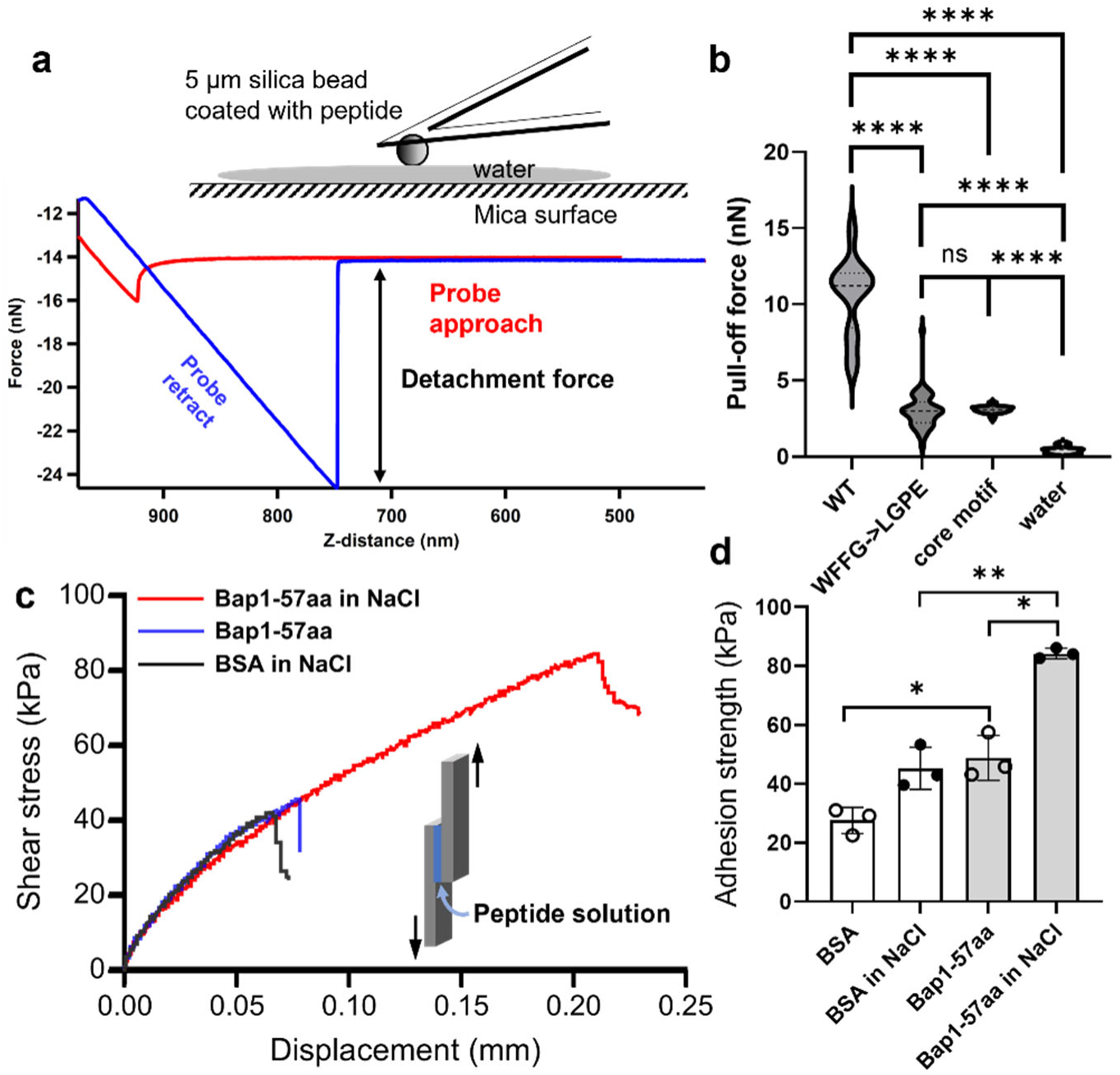
Atomic force microscopy (AFM) and lap shear test for characterizing the underwater adhesive strength provided by the biofilm-derived peptide. (**a**) A representative force-distance curve for the WT Bap1-57aa peptide in water. Inset: Schematic of the experimental setup. In brief, silica beads were first attached to the tip of AFM cantilevers, and subsequently coated with the peptide by incubating with 4 μM peptide in water. Next, the bead was allowed to approach a pristine mica surface in water and forces were measured. (**b**) Pull-off forces measured for the WT peptide, the WFFG→ LGPE mutant, the core motif, and the water control (without any peptide), presented as violin plots. Dashed lines correspond to 75th percentile (third quartile), median, and 25th percentile (first quartile). (**c**) Typical stress-extension curves of lap shear measurements of Bap1-57aa, in DI water and in 400 mM NaCl, and BSA in 400 mM NaCl, all prepared at 8 mg/mL. Inset: Schematic of lap shear measurements for two glass slides adhered by a peptide solution. (**d**) Measured adhesion strength of Bap1-57aa at 8 mg/mL, in comparison to the BSA control, in DI water and in 400 mM NaCl, respectively. All statistical analyses were performed using two-tailed *t*-tests with Welch’s correction. ns stands for nonsignificant (*p* > 0.05); * *p* ≤ 0.05; ** *p* ≤ 0.01; **** *p* ≤ 0.0001.

Using the Johnson-Kendal-Roberts (JKR) relationship (*56*, *57*), 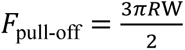, in which *R* is the bead radius (the nominal radius of curvature of the colloidal probes in this case) and W is the work of adhesion, we estimate W to be ≈ 1 mJ/m^2^. This value falls into the range of adhesive strengths measured for other underwater adhesive peptides/proteins (*58*). We also note that the procedure could be further optimized to increase W. For example, when varying the contact time after the beads come into contact with the mica substrates, we found that a short dwelling time (10 s in total) is helpful for proper adhesion; this could be related to the time needed for the peptide molecules to rearrange on the surface.

Complementing the microscale AFM measurements, we also performed lap shear measurements (*16*, *59*) to test the application potential of Bap1-57aa at the macroscopic scale. By sandwiching a Bap1-57aa solution at 8 mg/mL (≈ 1.2 mM) between two glass slides and briefly curing the sample, we measured the adhesion strength of the system while the sample was still wet. Typical stress-extension curves are shown in Fig. 5c and statistics are shown in Fig. 5d. We used BSA solutions at the same mass concentrations for comparison. We observed that using the particular preparation procedure (see Methods), we indeed obtained an appreciable adhesion strength (> 80 kPa) under wet conditions, particularly at salt concentrations comparable to oceanic water. The measured values are significantly higher than the BSA control and comparable to those measured for molecules designed based on Mfps at similar concentrations (*16*). Interestingly, we found that the measured strength is consistently higher in a salty solution than in DI water, suggesting that electrostatic screening promotes the adhesive functions of peptide. This may be related to the stronger aggregation tendency of the peptide when electrostatically screened, which in turn may be due to a competition between the interpeptide attraction caused by hydrophobic residues and repulsion between the positively charged residues.

### Potential application of the biofilm-derived peptide as a flocculant

During the fluorescence microscopy experiments performed in the bead-based adsorption assay, we observed that the Bap1-57aa peptide has a tendency to aggregate. Combining the aggregation tendency with the surface adsorption capability, we propose that the Bap1-57aa peptide may be used as a flocculating reagent to clear particulate contaminants from water. To demonstrate this point, we used well-dispersed suspensions of small (200 nm) PS particles as a model pollutant. Addition of FITC-labeled Bap1-57aa peptides to the particle suspension induced the formation of large co-aggregates with an interconnected network of beads and peptides, decreasing the number of particles remaining in solution (Fig. 6a). Interestingly, while the peptides can aggregate alone at the same concentration, the morphology of the co-aggregates is more compact – this suggests that while the peptide can bridge PS particles, the PS particles also help condense the protein aggregates. To generate mechanistic insights, we systematically increased the peptide concentration while keeping the bead concentration constant (Fig. 6b). At low peptide concentrations, most particles exist in free states (or settle on the glass substrate), whereas the FITC signals from the peptide colocalize precisely with the bead signal due to surface adsorption. As the peptide concentrations increase, the size and number of the co-aggregates increase while the number of free beads decreases (Fig. S7). At high peptide concentrations (> 1 μM), most particles are associated with the co-aggregates, demonstrating the efficacy of the biofilm-derived peptide as a flocculant for particle removal.

**Figure 6:**
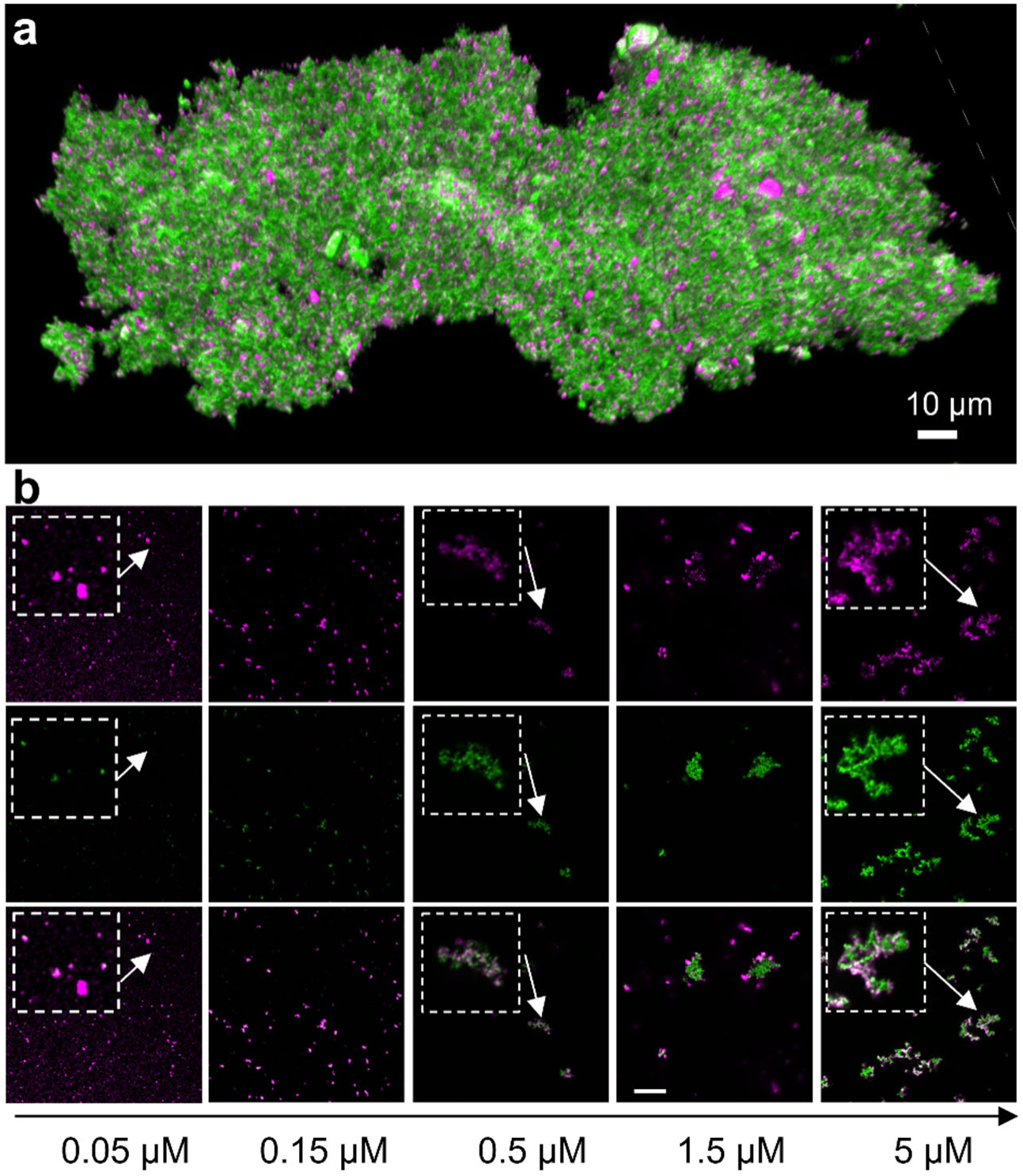
Bap1-57aa peptide as a flocculant. (**a**) A three-dimensional view of a large aggregate formed by the FITC-labeled Bap1-57aa (3 μM, green) and 200 nm PS beads (2×10^−4^ wt %, magenta) in M9 solution containing 0.2 mg/mL BSA. (**b**) Cross-sectional view of suspensions in PBS solution containing 200 nm PS beads (5×10^−3^ wt %, magenta) and increasing concentrations of FITC-labeled Bap1-57aa. Shown from top to bottom are the bead channel, the peptide channel, and the overlaid channel, respectively. Scale bar: 20 μm. Inset shows zoomed-in views for the highlighted region.

### Synthesis of the Bap1-57aa peptide from *E. coli*

While chemically synthesized peptides allowed us to perform quantitative assays, the cost will be prohibitive for large-scale use. As a proof-of-principle demonstration towards large-scale applications, we cloned the sequence into the pET-28b plasmid for *E. coli* expression and produced peptides as inclusion bodies using bacterial cultures. This plasmid introduces a poly-histidine tag at the N-terminus for purification while also allowing verification by Western blot. The recovery of the peptide from cell pellets, as well as the subsequent purification, turns out to be quite challenging, which is also commonly reported for adhesive peptides such as Mfps. Using the octaethylene glycol monododecyl ether (C_12_E_8_) detergent, inspired by the observation in QCM-D, we were able to solubilize the peptides in solution, as shown by Coomassie-stained SDS-PAGE gels and subsequent verification through Western blots (Fig. 7a). We were unsuccessful in further purification using a nickel column, likely due to peptide aggregation. Therefore, we used the crude inclusion body-produced peptide, which may also contain impurities including several larger proteins and DNA fragments, with the idea that a simpler purification step will be more suitable for large-scale applications down the road.

**Figure 7:**
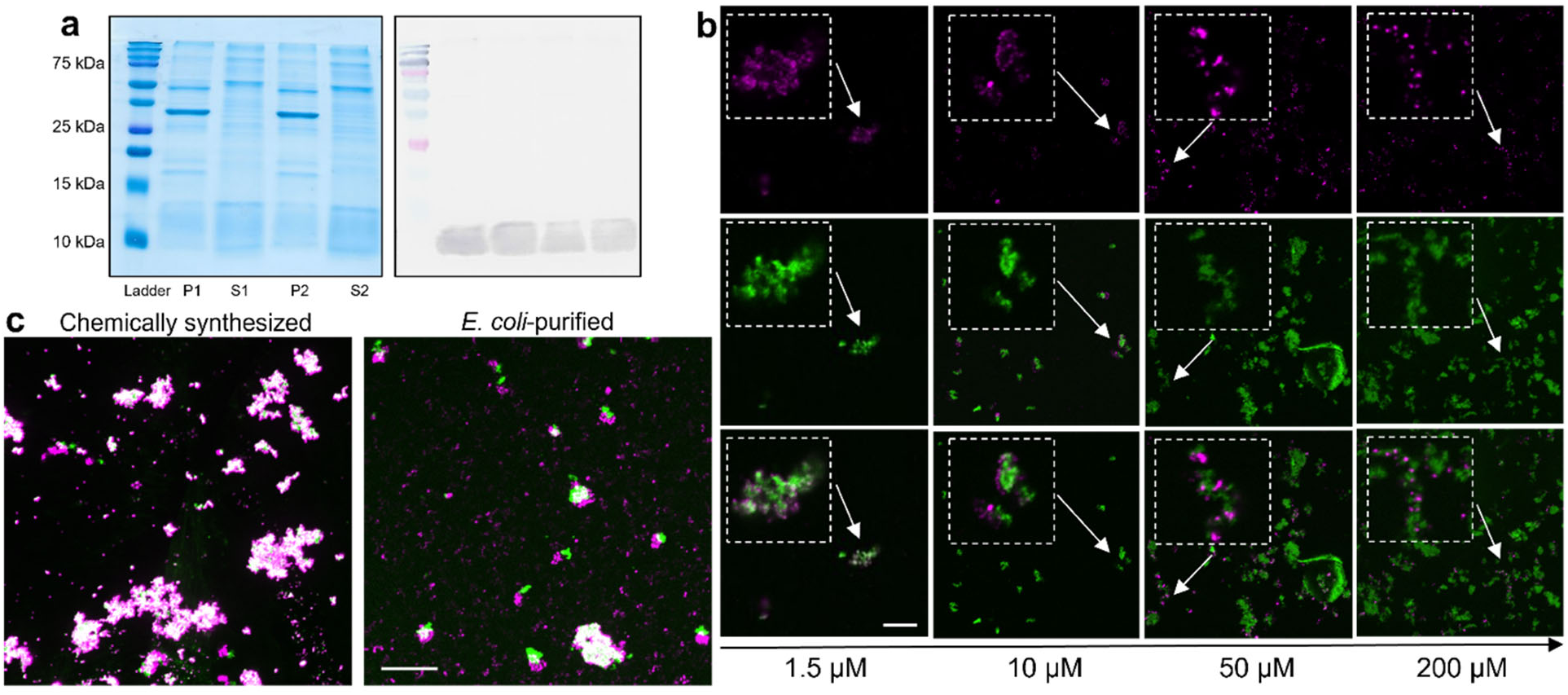
Bap1-57aa can be purified from *E. coli* in a functional form. (**a**) Coomassie stained gel (*left*) and the corresponding Western blot (*right*) showing expression and extraction of His-tagged Bap1-57aa peptide (≈ 9 kDa) from *E. coli*. P = pellet. S = supernatant extracted using C_12_E_8_ detergent. Two replicates (corresponding to the label 1 and 2) were shown to show reproducibility. (**b**) Cross-sectional view of suspensions containing 200 nm PS beads (5×10^−3^ wt %, magenta) and increasing concentrations of the *E. coli*-purified Bap1-57aa labeled with NanoOrange (green). Shown from the top to bottom are the bead channel, the peptide channel, and overlaid channel, respectively. Inset shows zoomed-in views for the highlighted region. (**c**) Maximum projection images (from the glass to 15 µm above) of co-aggregates of 200 nm PS beads with 5 µM chemically-synthesized (*left*) and *E. coli*-purified Bap1-57aa (*right*). Scale bars in b and c: 20 μm.

We repeated the co-aggregation assay with 200 nm PS beads and used NanoOrange to stain the peptide (Fig. 7b). Overall, we observed a similar ability to capture beads from the *E. coli*-purified peptide, indicating the success of the purification strategy. However, the concentration necessary for co-aggregation to happen is significantly higher than the chemically synthesized peptide. Specifically, the flocculating ability of the *E. coli*-purified peptide at 10 μM is comparable to chemically synthesized peptide at 0.5 μM or 1.5 μM. Mechanistically, the *E. coli*-purified peptides do form aggregates whose surfaces are covered by beads; however, we found many fewer PS beads inside the co-aggregates. As a result of the reduced effective surface area, at the same concentration (see for example 5 μM, Fig. 7c), the *E. coli*-purified sample has more free, uncaptured beads than the chemically purified peptides (Fig. S8), explaining the 5-20 fold difference in the efficiency of flocculation. Other factors such as inhibition of aggregation by the impurities or the C_12_E_8_ surfactant, the His-tag, or a combination of these factors are also possible. Future endeavors on the purification could further improve the performance of the peptide.

## Discussion

In this study, we employed a combination of characterization techniques including microscopy, spectroscopy, mechanical testing, and computer simulations to unravel the molecular mechanisms of surface association of a new biofilm-derived peptide. We find that the unique peptide originally discovered in the *Vibrio cholerae* biofilm adhesin Bap1 uses multiple sequence features to interact with a range of abiotic surfaces ranging from silica to polystyrene with various surface modifications. Using AFM and lap shear tests, we obtained a proof-of-concept demonstration that the peptide can be used to glue wet surfaces together both at the micron scale and at the macroscopic scale. This knowledge may open new avenues for developing bioinspired adhesives and biofilm control strategies. We have also successfully expressed and purified the peptide from *E. coli*, demonstrating that scalability is feasible, for potential future large-scale production.

The biofilm-derived peptide may offer several advantages over the popular design of bioadhesives inspired by Mfps. First, the absence of DOPA moieties means that it is chemically more stable and it can be applied in various aqueous environments. The simplicity in manipulating peptide sequences in bacteria also allows rapid testing and screening of optimized sequences for particular applications. Second, because of its bacterial origin and the absence of posttranslational modifications, it could be potentially mass-produced through fermentation, paving the way for scaling up its production for industrial use. Third, computational tools for gaining insights into molecular level information on how environmental factors impact conformation, folding, or aggregation, are readily available for peptides and proteins based on canonical amino acids. Lastly, the sequence can be easily integrated into other recombinant proteins to direct a protein to adhere to desired surfaces.

Previously, we have shown that the Bap1-57aa peptide also binds strongly to lipid bilayers (*34*, *44*), which is likely what the peptide has evolved for in the biofilm context. The ability to bind to both lipid membranes and abiotic surfaces renders Bap1-57aa peptide suitable as a multifunctional adhesive (*34*). In real-world environments, many abiotic surfaces are also contaminated by organic residuals including lipids and proteins and therefore creating a complex interface (*60*). Going forward, one could leverage the ability of the Bap1-57aa to interact with both lipids and abiotic substrates to design multifunctional underwater glues that achieve strong adhesion under diverse challenging conditions. Such adhesives could find widespread applications including binding wet surfaces, repairing tissues, stabilizing implants, or as priming agents to improve adhesion of subsequent protecting films for underwater infrastructure. Indeed, the peptide sequence has been successfully integrated into chitosan-based polymers for the use of mucoadhesives, recently (*61*). Finetuning of the peptide’s sequence or secondary structures, or integrating it into other functional domains could potentially broaden its utility as a versatile biomolecular tool.

Our previous work using surface force apparatus (SFA) has provided quantitative measurement of the adhesive strength the Bap1-57aa, as well as the core motif, between two mica surfaces (*58*). In that study, under comparable conditions, the pull-off force measured by the peptide based on the sequence of the core motif is similar to or even higher than that measured for Mfps, particularly Mfp5, the Mfp with highest DOPA content (*11*). Note that the pull-off force is affected by the ability of the peptide to adhere to surfaces, as well as the ability of the peptide to interact with other peptide molecules that defines the cohesive strength of the adhesive layer. Both adhesion and cohesion are affected by a range of environmental factors including ionic strength, nature of ions, pH, temperature, and surface properties including charge, hydrophobicity, and roughness (*62*). The different geometries, length scales, procedures, and rates at which the surfaces are displaced relative to each other in AFM and SFA may lead to the discrepancy in the results: in the SFA, prepared through a symmetric geometry, the core motif peptide generally performs better than the WT Bap1-57aa (*58*). In the AFM procedure presented here, the peptide molecules need to coat the beads surface first, which may explain why the WT Bap1-57aa performs the best considering its stronger ability to adsorb onto the silica surface. Further work characterizing the film thickness, molecular arrangement and hysteresis, and morphology of the adhesive film may shed light on this difference and more importantly, the failure mode during pull off.

Intrinsically disordered peptides or proteins (IDPs) have received significant attention recently due to their ability to form membraneless organelles or condensates in cells (*63–66*). However, IDPs can also display transient secondary structures when interacting with other molecules (*65*, *67*). Indeed, we found that Bap1-57aa displays different conformations when contacting different surfaces. Previously, using a combination of circular dichroism, fluorescence spectroscopy, and full-atom simulations (*44*), we have shown that the core motif (SYWFFGWHTK) forms a β-hairpin to insert into lipid membranes, while the peripheral repeating units increase lipid binding through avidity. On the other hand, the full Bap1-57aa peptide assumes a mixture of β-strands and random coils in solution, and only turns into the β-hairpin conformation upon contacting lipids. Here, combining SFG measurement and MD simulations, we observed yet another conformation (largely α-helix) when adsorbing on negatively charged silica surfaces. The flat energy landscape of IDPs allows facile conversion between conformations, which can be heavily influenced by an external surface. This principle could potentially be applied to the future design of peptide-based adhesives.

The ability to form co-aggregates with particles not only enables the use of the biofilm-derived peptide as flocculant, but also further opens the possibility of modifying its adhesive properties with particulate inclusions (*68*). In general, aggregation can be harnessed as a way for the mass production of adhesives, with potential to control aggregate structure, function, and size, by using solvent conditions as variables to tune dissipation pathways and to amplify the toughness of protein/peptide-based adhesives. For example, recently, Wilson *et al.* chemically denatured BSA using urea to cause aggregation and developed an injection method for the mass production of underwater glues (*68*). Even cheaper proteins such as Zein (*59*) and soy proteins (*69*) have also been employed with various denaturing methods for gluing wet surfaces such as water-soaked wood. Along this line, the ability to produce the Bap1-57aa peptide from bacterial culture is critical for future applications on a large scale. Looking forward, the mechanism underlying the aggregation tendency of the Bap1-57aa, as well as the contribution of the aggregation to adhesive function, warrants further mechanistic studies.

## Materials and Methods

### Source of materials

Chemically synthesized peptides are purchased from either Atlantic peptides or LifeTein. Results from the two sources are generally consistent. 5 µm Polystyrene beads were purchased from Invitrogen (Nonionic #N37460; Sulfate #S37227; Carboxylate #C37255). Silica beads were purchased from Polysciences (Silica #25348; Silica-NH_2_ #24758).

### Bacterial strains

All *V. cholerae* strains used in this study were derivatives of the wild-type *V. cholerae* O1 biovar El Tor strain C6706str2 and listed in Supplementary Table 3. A rugose strain background was used, which harbors a missense mutation in the *vpvC* gene (*vpvC*^W240R^) that elevates intracellular c-di-GMP levels (*70*). The rugose strains form robust biofilms and thus allow us to focus on the biochemical mechanisms governing surface binding rather than mechanisms involving gene regulation. For the biofilm adhesion assay, the other adhesin RbmC was further deleted to avoid confounding factors (*34*). For the matrix staining experiment, the functional copy of RbmC is present; otherwise nonadhesive biofilms will be washed away during the washing step. Moreover, all biofilm assays are performed in a Bap1 background in which the β-prism is deleted; previous work has shown that the cap of the β-prism contains several exposed lysine and tryptophan residues that confound the interpretation of the biofilm adhesion results (*34*). Additional mutations were genetically engineered into this *V. cholerae* strain using the natural transformation (MuGENT) method (*71*).

### Bacterial growth

All strains were grown overnight in lysogenic broth (LB) at 37 °C with shaking. 1× M9 salts were filter sterilized and supplemented with 2 mM MgSO_4_ and 100 µM CaCl_2_ (abbreviated as M9 medium below). Biofilm growth was generally performed in M9 medium supplemented with 0.5 % glucose.

### Bead-based adhesion assay

First, the surface of the 96-well plate was treated with NaOH to render it more hydrophilic and negatively charged. Briefly, before adding the solution, 100 µL of 10M NaOH aqueous solution was added to the wells and incubated at room temperature for 10 min, after which the wells were washed with DI water until the pH was neutral. Chemically synthesized fluorescently labeled peptides (Atlantic or Lifetein peptides) were dissolved and stored in DMSO at 150 µM. 100 µL of water containing 1 mg/mL BSA, various concentrations of FITC-labeled peptide or dextran-FITC and 0.01 % (weight percent) 5 µm microspheres was shaken in Eppendorf tubes for 1 h at room temperature. The sample was then spun down and resuspended in 10 mM Tris buffer pH 7.4 and 150 mM NaCl before transferring to a NaOH-treated 96-well plate with a glass bottom (MatTek P96G-1.5-5-F) and allowed to settle at room temperature for 5 min before imaging. Thus-prepared samples were imaged with a spinning disk confocal microscope (Nikon Ti2-E connected to Yokogawa W1) using a 60× oil objective (numerical aperture = 1.40) and a 488 nm laser excitation and bright field. For each sample, at least three locations were imaged and captured with an sCMOS camera (Photometrics Prime BSI). Each field of view contained roughly 100–150 beads.

### Quantification of bead adsorption assay

The background signal due to the camera in the 488 nm channel was measured by taking images of M9 medium and quantifying them with built-in functions of the Nikon Element software. After subtracting the background signal, the signal intensity per unit area on the surface of the beads and in the solution was calculated using custom MATLAB codes and the difference was determined to give the excess surface signal.

### Quartz crystal microbalance

5 MHz 14 mm wrapped silicon oxide sensors from AWSensors (AWS SNS 000049 A) were used fresh from the package within one week of opening the package. No additional cleaning was performed, as directed by the supplier. Frequency at the overtones 15, 25, 35, and 45 MHz was monitored using the OpenQCM Next device. Data reported in Figure 2 and in the main text are from the 5^th^ (25 MHz) harmonic, although the results were the same within error across all overtones. Dissipation was also monitored at all overtones and remained low (<5 ppm difference between water and water with the peptide film present) for the duration of the experiment, indicating mass can be calculated using the linear Sauerbrey relationship with frequency change. Peptide solutions were prepared by diluting 150 µM stock in DMSO into ultra-pure water. For 1 µM peptide, the final DMSO concentration was ≈ 0.7 % (volume fraction). This amount of DMSO was found to change the frequency compared to water, so values reported in Figure 2 are in pure water, since little to no peptide washed off with each water injection. Water, isopropanol, and 0.7 % DMSO was injected into the device prior to adding peptide to rinse the system and measure baseline values for water. In one experiment, the chip was washed with water, ethanol, and isopropanol before measuring the value of two water injections for 40 seconds each. After 1 µM peptide was injected, the chip was washed with subsequent water, 50 mM Tris and 100 mM KCl, water, isopropanol, and again water. Results obtained are similar to the other three data sets.

### Biofilm adhesion assay

Overnight cultures of the indicated strains constitutively expressing mNeonGreen were grown from individual colonies at 37 °C with shaking in 1.5 mL LB. 50 µL from each culture was used to inoculate 1.5 mL of M9 medium supplemented with 0.5 % glucose and grown at 30 °C with shaking until the OD_600_ was between 0.1 and 0.3. The cultures were then diluted to an OD_600_ ≅ 0.001. 100 μL of the regrown culture was aliquoted into the wells of a 96-well plate with a glass bottom (MatTek P96G-1.5-5-F) and incubated at 30 °C for 1 hour. The wells were then washed twice with M9 medium and replaced with M9 medium with 0.5 % glucose and 0, 0.2, 0.4, 0.6, 0.8, 1, or 1.2 mg/mL BSA. The lid was secured with a layer of parafilm and the 96-well plate was subsequently incubated at 30 °C for 16 hours. Thus-prepared samples were imaged with a spinning disk confocal microscope (Nikon Ti2-E connected to Yokogawa W1) using a 60× water objective (numerical aperture = 1.20) and a 488 nm laser excitation. For each sample, several locations with 3×3 tiles where imaged and captured with an sCMOS camera (Photometrics Prime BSI). The *x*-*y* pixel size was 0.22 μm and the *z*-step size was 3 μm. The wells were then washed twice with M9 medium and re-imaged at the same locations. All images presented in this study are raw data rendered using the Nikon Elements software.

### Quantification of biofilm adhesion assays

Image analysis was performed with built-in functions of the Nikon Elements software by thresholding each image layer-by-layer and measuring the total binarized area above the threshold in each layer. The binary area for each sample *z*-slice was then summed to give the total biovolume, and the ratio of the total biovolume after versus before the washing step was calculated.

### *In situ* biofilm immunostaining

Overnight cultures of the indicated strains with *bap1* tagged with 3×FLAG at the C-terminal and constitutively expressing mNeonGreen were grown following the same procedure as described above. The initial incubation time was adjusted to 30 minutes when biofilms were grown in the presence of BSA. The wells were washed twice with M9 medium; subsequently, 100 µL of M9 medium with 0.5 % glucose, 0.5 mg/mL BSA (Sigma-Aldrich A9647), and 2 µg/mL anti-FLAG antibody conjugated to Cy3 (Sigma-Aldrich A9594) was added to the well. The lid was secured with a layer of parafilm and incubated at 30 °C for 40 hours. Thus-prepared samples were imaged with a spinning disk confocal microscope (Nikon Ti2-E connected to Yokogawa W1) using a 100× oil immersion objective (numerical aperture = 1.35) or a 60× water immersion objective (numerical aperture = 1.20) and a 488 nm laser excitation to observe the cells and a 561 nm laser excitation to observe protein localization, with the corresponding filters. The images were captured with an sCMOS camera (Photometrics Prime BSI) at a *z*-step size of 0.5 µm.

### Molecular dynamics simulations

To examine the binding behavior of Bap1-57aa on an abiotic silica surface, silica (a-Quartz) slab with dimension of 100 Å × 100 Å × 10 Å was constructed at neutral pH = 7 using *nanomaterial modeler* in the CHARMM-GUI web server (*72*) and solvated with a water box of 100 Å × 100 Å × 160 Å. A disordered structure of the Bap1 peptide (Tyr415-Lys471; 57 amino acids) was generated using TraDES (*73*) and placed near the silica surface. Overlapping water molecules within 1.5 Å of the peptide in the combined system (peptide + silica slab) were removed to avoid steric clashes. Na^+^ and Cl^−^ ions were added to neutralize the system and the ionic strength of the system was maintained to 150 mM NaCl. The total number of atoms was 156124, including 47567 water molecules. Silica force fields integrated into CHARMM was used for the SiO_2_ nanomaterials (*74*). CHARMM36m (*75*) and TIP3P water model (*76*) force field were used for protein and water, respectively.

All simulations were performed using NAMD 3.0 (*77*). System was energy minimized for 10,000 cycles using the conjugate gradient and subsequently equilibrated at constant NVT (constant temperature and volume) for 250 ps with a 2 fs timestep. Positional restraints were applied to the heavy atoms of silica surface, while harmonic restraints were imposed on the protein backbone with a force constant of 10 kcal mol^−1^ Å^−2^. Three successive equilibrations at constant NPT (constant temperature and pressure) simulations were performed for 1, 4, 20 ns with 2 fs timestep, during which a force constant on the protein backbone was gradually reduced to 5, 2 and 1 kcal mol^−1^Å^−2^, respectively. Finally, production simulations were performed for 0.6 µs in 10 replicates at constant NPT. Snapshots were saved at 100 ps interval and the last 500 ns of each trajectory were considered for further analysis.

Long-range electrostatic interactions were treated using the particle mesh Ewald (PME) method (*78*). All bonds involving hydrogen atoms were constrained with the SHAKE algorithm (*79*). Nonbonded interaction cutoff was set to 12 Å cutoff with force switching starting at 10 Å. The temperature was maintained at 300 K using a Langevin thermostat with a damping coefficient of 1 ps^−1^, and the pressure was controlled at 1 atm using the Nosé-Hoover Langevin piston method (*80*).

### Analysis of simulation data

CPPTRAJ (*81*) and in-house python scripts were used to calculate averaged secondary structure frequency for 10 replicate simulations. Contact frequency within 3.5 Å cutoff between heavy atoms and silica surface was calculated using tcl script in VMD (*82*) for each simulation and averaged over replicate simulations.

### Sample preparation for SFG experiments

The Bap1-57aa peptide was added to pure water at 20 µM and sonicated for two intervals of 3 minutes for dissolution. Then, 20 µL of the sonicated solution was added onto a clean glass surface and allowed to dry overnight in a desiccator to form a film for the SFG experiments. Two other samples were prepared using the same procedures by replacing H_2_O with D_2_O or 40 % DMSO as the solvent.

### SFG spectrometer setup and data acquisition

A home-built broad bandwidth SFG spectrometer was used to collect SFG spectra with the *ssp* polarization (*s*-polarized SFG, *s*-polarized visible and *p*-polarized IR beams). The experimental setup and the protocols used to collect and process SFG data were described in detail in previous work (*83*, *84*). In this study, SFG spectra were recorded for 10 minutes. The IR beam was tuned to the vibrational regions corresponding to the amide I and NH stretching regions for probing protein secondary structures. The resulting spectra were fitted into the Lorentzian line shape:

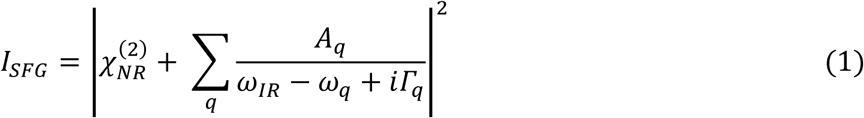

where *I*_SFG_ is the SFG intensity; 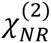 the non-resonant contribution of the second order susceptibility; *ω*_IR_ the incident IR frequency; *A*_q_ the amplitude; *ω*_q_ the resonant frequency; Г_q_ the damping factor for the *q*^th^ vibrational mode. Error bars for the fitted parameters were computed in the data analysis software Igor Pro (version 9.01), with each error bar representing ±1 standard deviation estimated from the fit residuals and covariance matrix, assuming normally distributed residuals.

### Atomic force microscopy

AFM probes for adhesion measurement were prepared as described previously (*40*, *85*). Silica microspheres (Polysciences, Warrington, PA) of diameter 5.0 µm were attached to tipless AFM cantilevers (PNP-TR-TL-Au, NanoWorld, Switzerland) using UV curable optical adhesive (NOA 61, Norland Products). The probes were immersed in water containing 4 µM peptide for 20 min, then replaced with Milli-Q water. Force spectroscopy was performed with Asylum Cypher ES atomic force microscope. Probe calibration and spring constant determination were achieved via the thermal noise method. Curves were generated using a constant approach and retraction velocity of 0.5 µm/s. A retraction was triggered upon applying a 1 nN force following a 5 s dwell on the surface. Each force curve was analyzed individually in Igor Pro software. The adhesion force was taken to be the difference between the minimum and baseline forces in the retraction curve. Force curves that did not show a clear adhesion event were excluded from the analysis. 42 force curves for WT, 42 force curves for core motif, 46 force curves for WFFG->LGPE, and 27 force curves for water only control were obtained for analysis.

### Lap shear test

In the lap shear experiments, hydrophilic glass surfaces were utilized as the substrates to evaluate the adhesion performance of chemically synthesized Bap1-57aa. The glass slides were cleaned by sonication in acetone for 10 min, followed by extensive rinsing with water and drying under a nitrogen stream. To prepare the specimens, 5 μL of the peptide/protein solution (Bap1-57aa or BSA) at a concentration of 8 mg/mL was applied onto a predefined 1 cm × 1 cm area at the center of one glass substrate. The peptide/protein was dissolved in either DI water or a 400 mM NaCl solution. The solution was evenly spread across the 1 cm² bonding area using a pipette tip. A second glass slide was then overlapped to form the lap joint. During assembly, moderate rubbing and pressing were applied manually to ensure the formation of a uniform, thin film and to eliminate any trapped air bubbles. The assembled specimens were allowed to equilibrate for 3 min prior to testing: at room temperature for samples in 400 mM NaCl solutions, and at 55 °C for samples in DI water. All lap shear tests were performed using an Anton Paar MCR Rheometer (502WESP) equipped with a highly sensitive normal force (F_N_) sensor. The bonded specimens were mounted on the rheometer stage using the accessory #129538 (solid torsion bar fixture SRF12/CTD V2.0), and the adhesion strength was measured in a pull-off mode with a constant separation speed of 150 μm/s. The adhesion strength was determined by dividing the peak detachment force (the absolute value of the maximum F_N_ recorded) by the bonded overlap area (1.0 × 10^−4^ m^2^). Each condition was tested with at least three independent replicates, and the results are expressed as mean ± standard deviation (SD).

### Bap1-57aa expression and purification from *E. coli*

Bap1-57aa peptide expression and purification was performed in a similar manner as previously described for Bap1 protein (*86*). DNA sequence of the 57aa, located within the β-prism domain of Bap1, was cut out and inserted into pET28b(+), using NdeI and XhoI restriction sites to generate a N-terminally His-tagged construct containing a thrombin cleavage site. The resulting sequence is: MGHHHHHHSSGLVPRGSHMDYLGLEWKTKTVPYLGVEWRTKTVSYWFFGWHTKQVAYLAPVWKEKTIPYAVPVTLSK. This construct was transformed into T7 Express *E. coli* (New England Biolabs), and overnight cultures were diluted 60-fold into fresh LB-Miller media (MP Biomedicals) supplemented with 50 µg/mL kanamycin monosulfate (GoldBio). Cultures were incubated at 37 °C with shaking at 200 rpm until reaching an OD_600_ between 0.5 and 0.8. Protein expression was induced by and addition of 1 mM isopropyl its β-D-1-thiogalactopyranoside (IPTG) (GoldBio), followed by an additional incubation for 4 hours at 37 °C with shaking at 200 rpm. Cells were pelleted by centrifugation in a Sorval LYNX 6000 centrifuge using an F9-6X-1000 LEX rotor at 5422 × g for 15 min at 4 °C. Cell pellets were resuspended in 1× TBS (20 mM Tris, pH = 7.5, 150 mM NaCl) to a final volume of ≈ 10 mL per pellet from 1 liter of expression culture. Samples were stored at −80 °C until further processing. Thawed cell suspensions were supplemented with 1 mM protease inhibitor 4-(2-Aminoethyl)benzenesulfonyl fluoride hydrochloride (AEBSF) (GoldBio) and lysed via passage 5 times through an EmulsiFlex-C5 high pressure homogenizer (Avestin, Inc.) at ≈18,000 psi (124.1 MPa). The cell lysate was divided into two equal batches that were simultaneously centrifuged in a Sorvall LYNX 6000 at 21734 × g for 30 min using an F20-12X50 rotor and further purified.

After centrifugation, the supernatant was discarded by decantation and the pellet was resuspended by vortexing in 5 mL of Detergent Wash Buffer (2× Critical Micelle Concentration (CMC) of 3-[(3-Cholamidopropyl)-dimethylammonio]-2-hydroxy-1-propane sulfonate (CHAPSO) in 1× TBS) per pellet from 1 liter of expression culture. Resuspended pellets were stirred on a magnetic stirrer on ice for 30 min and centrifuged in a Sorvall LYNX 6000 at 21,734 × g for 30 min using an F20-12X50 rotor. Pellet resuspension was repeated with four different buffers: High Salt Wash Buffer (1 mol/L (M) NaCl, 1× TBS), DNAse Wash Buffer (10 mM CaCl_2_ and up to ≈100 µg/mL DNAse I), 1× TBS Wash Buffer (20 mM Tris, pH = 7.5, 150 mM NaCl) and Solubilization Buffer (4× CMC C_12_E_8_ (Octaethylene Glycol Monododecyl Ether, 1× TBS), respectively. All buffers were filtered using a 0.22 µm filter. Pellets resuspended in the Solubilization Buffer were subsequently stirred at room temperature. The soluble fraction (supernatant) containing Bap1-57aa peptide was stored at room temperature and the insoluble fraction (pellet) was stored at 4 °C.

### Gel electrophoresis (SDS-PAGE)

Samples were analyzed by SDS-PAGE using gels prepared with 15 % bisacrylamide (Bio-Rad), 25 mM Tris, pH = 8.3, 0.05 % ammonium persulfate (Thermo Fisher Scientific), and 0.025 % TEMED (Bio-Rad). Electrophoresis was conducted at 180 V for 1 h at room temperature in a running buffer composed of 25 mM Tris, 192 mM glycine, 1 % SDS, pH = 8.35. Following electrophoresis, gels were stained overnight with 0.02 % Coomassie Brilliant Blue G-250 in 10 % acetic acid, and subsequently destained in 10 % acetic acid, using Precision Plus Unstained Protein Standards (BioRad) as the ladder. In another set of experiments, gels were used for transfer to PVDF membrane; in these cases, Precision Plus Dual Color Protein Standards (BioRad) were used as the ladder.

### Western Blot

Protein transfer to PVDF membrane was carried out at 100 V for 1 h with an ice bag on the magnetic stirrer. Transfer buffer consisted of 25 mM Tris, 192 mM glycine, pH = 8.3 and was incubated at −20 °C for 20 min. After the transfer, the PVDF membrane was washed 4 times with 1× PBS-T buffer (137 mM NaCl, 2.68 mM KCl, 10.15 mM Na_2_HPO_4_, 1.76 mM KH_2_PO_4_, 0.05 % (volume fraction) Polyoxyethylenesorbitan, Monolaurate (Tween 20), pH = 7.4) at 5-min intervals with shaking at room temperature. The membrane was subsequently blocked with lyophilized milk dissolved in 1× PBS-T buffer for 2 h with shaking at room temperature, followed by 3 times of washing with 1× PBS-T buffer at 5-min intervals with shaking at room temperature. Next, the membrane was incubated overnight with Goat Ani-Mouse IgG (H+L-HRP Conjugate) (BioRad) primary antibodies with shaking at 4 °C, followed by 3 times of washing with 1× PBS-T buffer at 5-min intervals with shaking at room temperature. Subsequently, the membrane was incubated with 6×His Tag Monoclonal Antibody (HIS.H8, Invitrogen) secondary antibodies for 1 h with shaking at room temperature, followed by 3 times of washing with 1× PBS-T buffer at 5-min intervals with shaking at room temperature. Lastly, the membrane was developed using Opti-4CN Substrate Kit (BioRad), followed by washing in deionised autoclaved water for 15 min at room temperature without shaking.

### Co-aggregation assay

96-well glass plate was treated with NaOH by adding 100 µL of 10 M NaOH to each well and incubating for 10 min. After the incubation, the NaOH solution was removed and each well was washed 6 times with 200 µL of DI water, and the plate was left dry. Chemically synthesized Bap1-57aa peptide labeled with FITC was dissolved in DMSO and diluted to concentrations of 0.015 µM, 0.05 µM, 0.15 µM, 0.5 µM, 1.5 µM, and 5 µM. To each sample, 200 nm FluoroSpheres carboxylate-modified beads (Invitrogen # F8807) was added to a final concentration of 0.005 % in 1× PBS buffer in a final volume of 50 µL. Negative controls with beads only and peptide only (at a concentration of 1.5 µM) were also prepared. Each sample was briefly vortexed, covered with aluminium foil and incubated on a shaker for 1 h. After the incubation, thus-prepared samples were transferred to the 96-well plate and imaged with a spinning disk confocal microscope (Nikon Ti2-E connected to Yokogawa W1) using a 60× oil objective (numerical aperture = 1.40). A 488 nm laser excitation was used to image the peptide, and a 640 nm laser excitation was used to image the polystyrene beads, with the corresponding filters. For each sample, at least six locations were imaged and captured with an sCMOS camera (Photometrics Prime BSI), from which a representative image was shown for each condition in the figures.

Bap1-57aa peptide purified from *E. coli* using Solubilization Buffer (4× CMC C_12_E_8_ in 1× TBS) was diluted to concentrations of 1.5 µM, 5 µM, 10 µM, 25 µM, 50 µM, 100 µM, 200 µM, and 400 µM. To each sample, 200 nm FluoroSpheres carboxylate-modified beads (Invitrogen # F8807) was added to a final concentration of 0.005% in 1× PBS to a final volume of 50 µL. Also added to the solution was 50× diluted NanoOrange (Thermo Fisher Scientific N6666) to stain the peptide. Negative controls with beads only and peptide only (at a concentration of 50 µM, with and without NanoOrange) were also prepared. Each sample was briefly vortexed, covered with aluminium foil and incubated on a shaker for 1 h. After the incubation, thus-prepared samples were transferred to the 96-well plate and subsequently imaged using the same setting as above.

### Statistics and reproducibility

Error bars correspond to standard deviations from measurements taken from distinct samples unless indicated otherwise. Standard *t*-tests were used to compare treatment groups and are indicated in each figure caption. All statistical analyses were performed using GraphPad Prism software. Microscopy images and spectra were shown from representative results from at least three independent experiments.

## Supporting information

All supplementary figures and tables combined

## Data availability

All final data are available in the main text or the supplementary materials.

## Materials availability

All bacterial strains constructed as part of this work will be provided to the community upon request in a timely fashion and shipped in accordance with biosafety standards and regulations.

## Acknowledgements

We thank Dr. Caitlin Davis and Ms. Sydney Shuster for helpful discussions.

## Author Contributions

X.H. and J.Y. conceptualized the project. X.H. performed strain construction and validation. X.H. performed bead-based assays and quantification. X.H. performed adsorption assays and biofilm imaging. M.E.M. performed QCM-D measurement and data analysis. X.H. and P.D. performed the AFM measurement. R.P. and H.-X.Z. performed MD simulations and analyzed MD data. Y.L. performed lap shear measurements. E.L. and M.W. purified the peptide from *E. coli*. E.L. and S.S. performed the flocculation assay. Z.W., R.A.J., and E.C.Y.Y. performed the SFG measurements and data analysis. X.H., R.P., Z.W., M.E.M., E.C.Y.Y., H.-X.Z and J.Y. wrote the manuscript. All authors contributed to the final manuscript.

## Data availability

All data needed to evaluate the conclusions in the paper are present in the paper and/or the Supplementary Materials.

## List of material contained in the Supplementary Material

Supplementary Figures 1-8, Supplementary Table 1-3, and References.

