## Supplementary material for "A biofilm-derived peptide as an underwater adhesive": All supplementary figures and tables combined

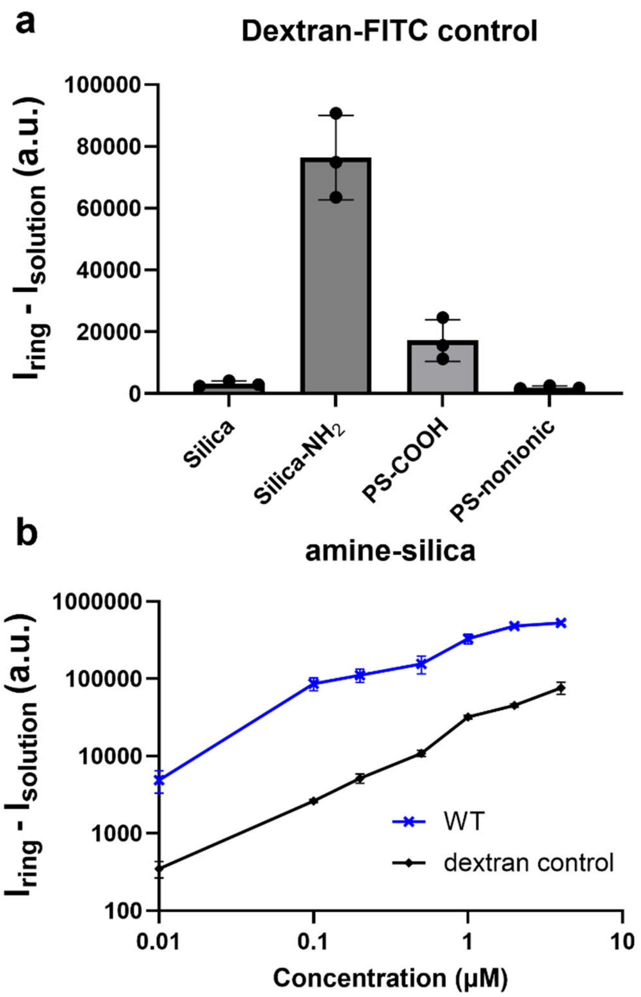


**Fig. S1**: **The amine-functionalized silica beads show enhanced nonspecific adhesion in the fluorescence-based adsorption assay.** (**a**) Quantification of excess signals of the negative control (FITC-labeled dextran at 4 μM) on different microspheres. (**b**) Adsorption curves of FITC-labeled Bap1-57aa and dextran control on amine-functionalized silica spheres.

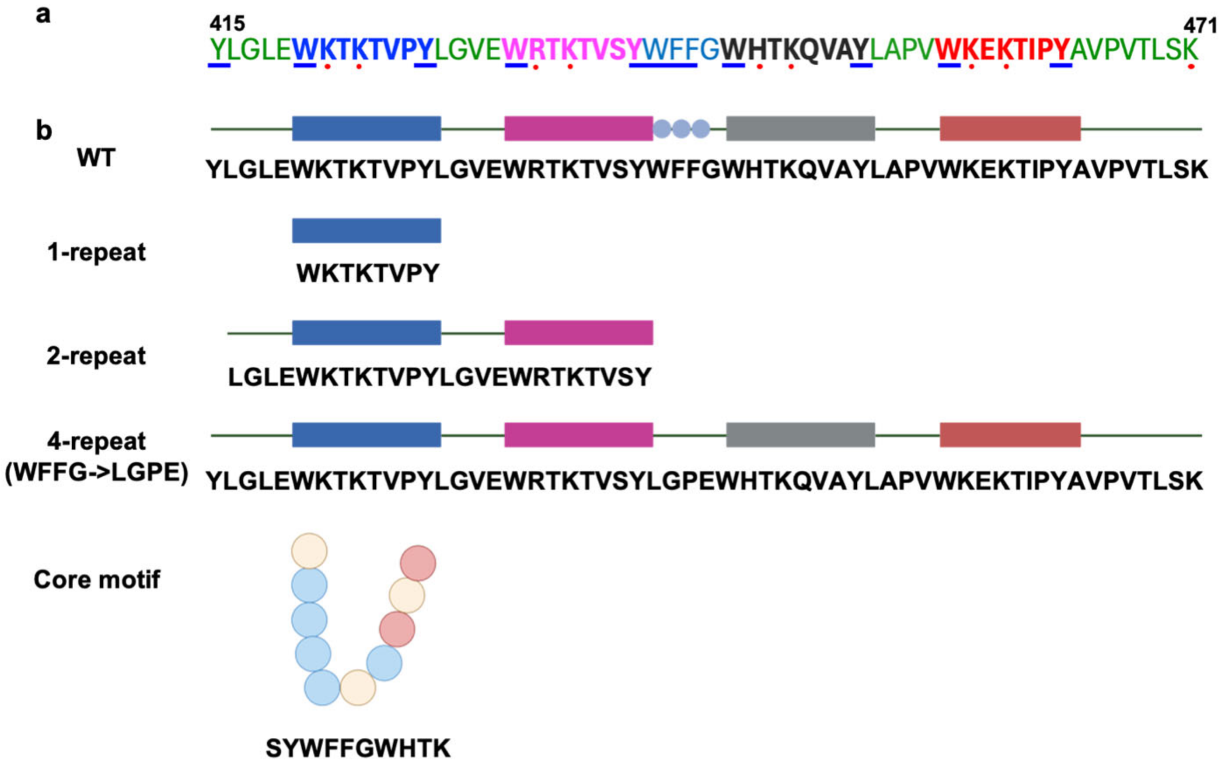


**Fig. S2**: **Schematics and sequences of different peptides used in this study.** (**a**) Sequence of WT Bap1-57aa. Aromatic and basic residues are indicated by blue underline and red dots, respectively. Four pseudo repeats are shown in blue, magenta, black, and red. (**b**) Schematic representation and sequences of various peptides in this study. Colored circles indicate amino-acid types: blue, aromatic; red, positively charged; beige, other.

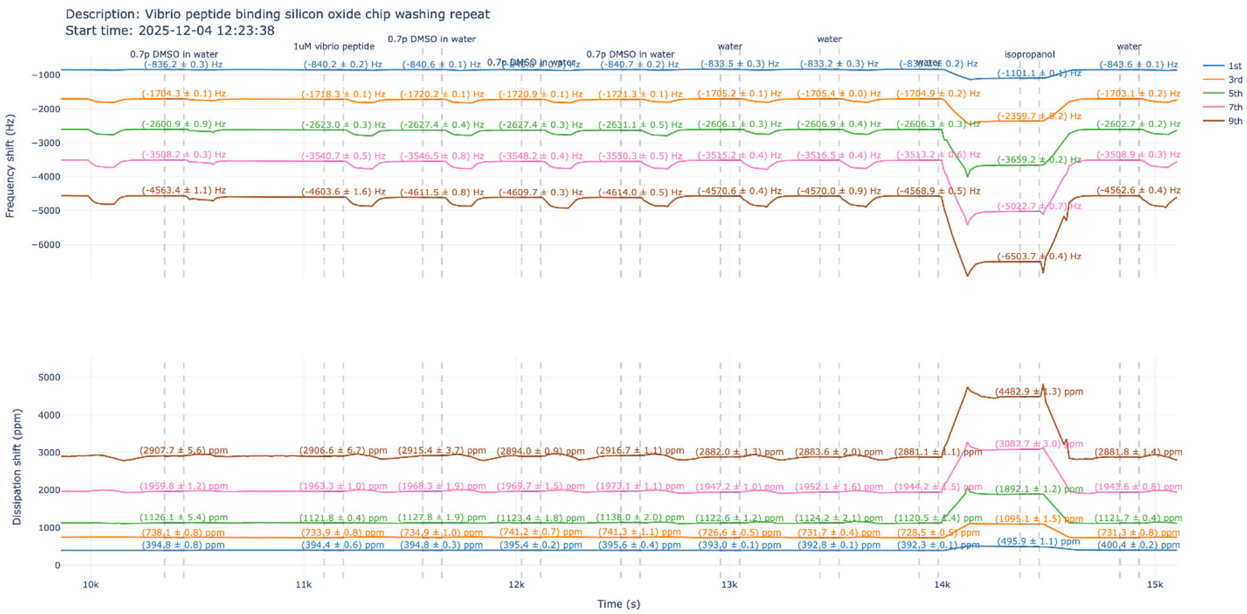


**Fig. S3**: **Raw QCM-D data.** Shown are raw QCM-D data corresponding to the plot in Figure 2a, showing all odd overtones from the 1$^{st}$ to 9$^{th}$.

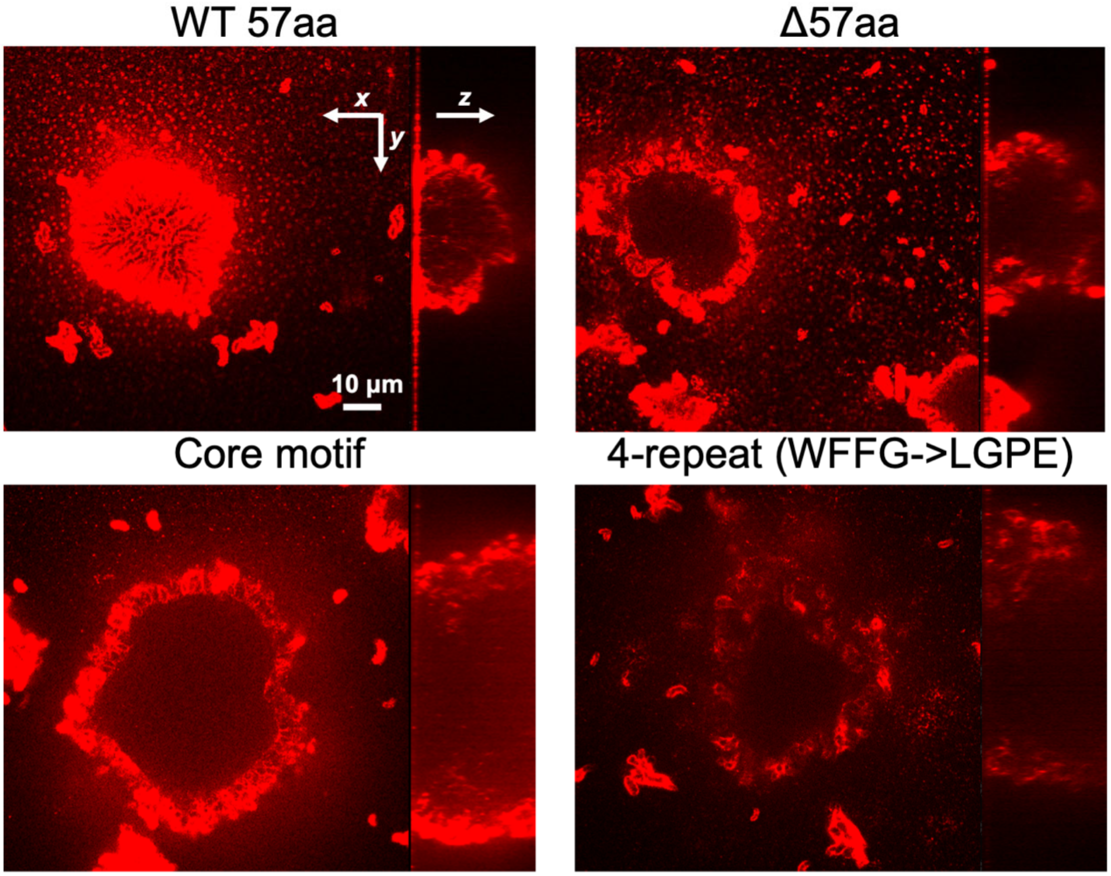


**Fig. S4**: **Localization of 57aa in *V. cholerae* biofilms.** Biofilms formed by different mutants possessing different 57aa variants, tagged by 3×FLAG at the C-terminus, were stained using an Anti-FLAG antibody conjugated to Cy3 (red). 0.5 mg/mL of BSA was used to block nonspecific surface binding. Shown are cross-sectional images at the glass substratum as well as the side views.

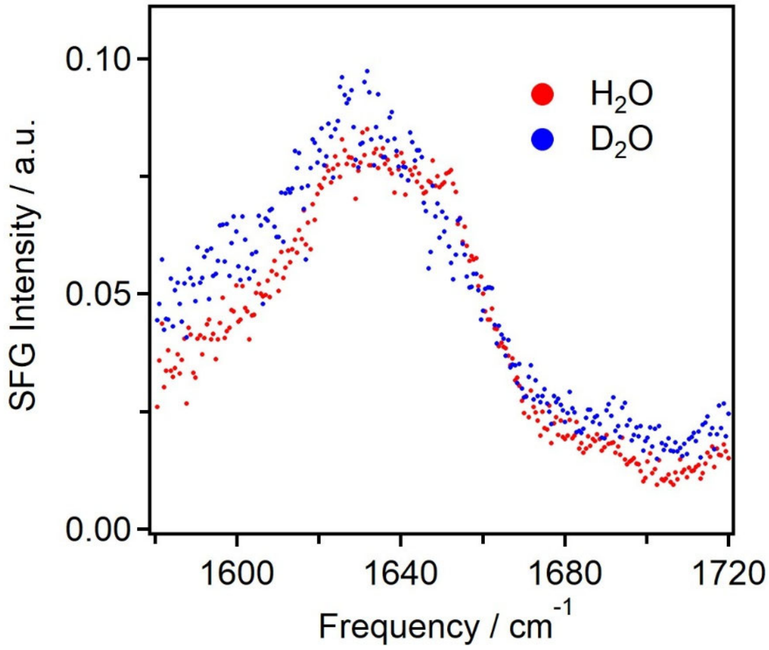


**Fig. S5**: **Additional SFG data.** Shown are sum frequency generation spectra in the amide I region of the Bap1-57aa peptide deposited on a glass slide prepared using $H_2O$ (red) or $D_2O$ (blue) as solvents.

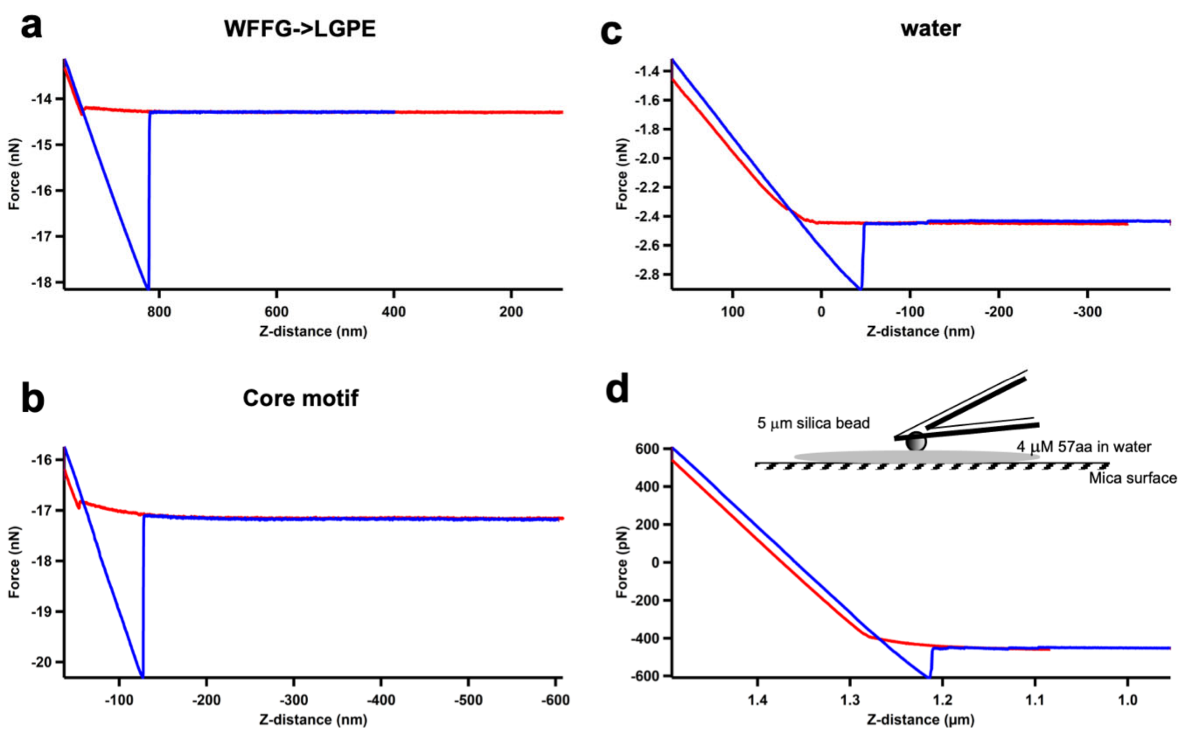


**Fig. S6**: **Additional AFM results.** (**a-c**) Representative force-distance curves for the WFFG->LGPE variant (a), the core motif (b), and the water control (c). (**d**) Alternative AFM measurement geometry. We have also tried another configuration in which the peptides are present in the aqueous solution so that both the mica surface and bead surface are coated with the peptide (Inset); however, we observed a much lower adhesive force and inconsistency between different measurements.

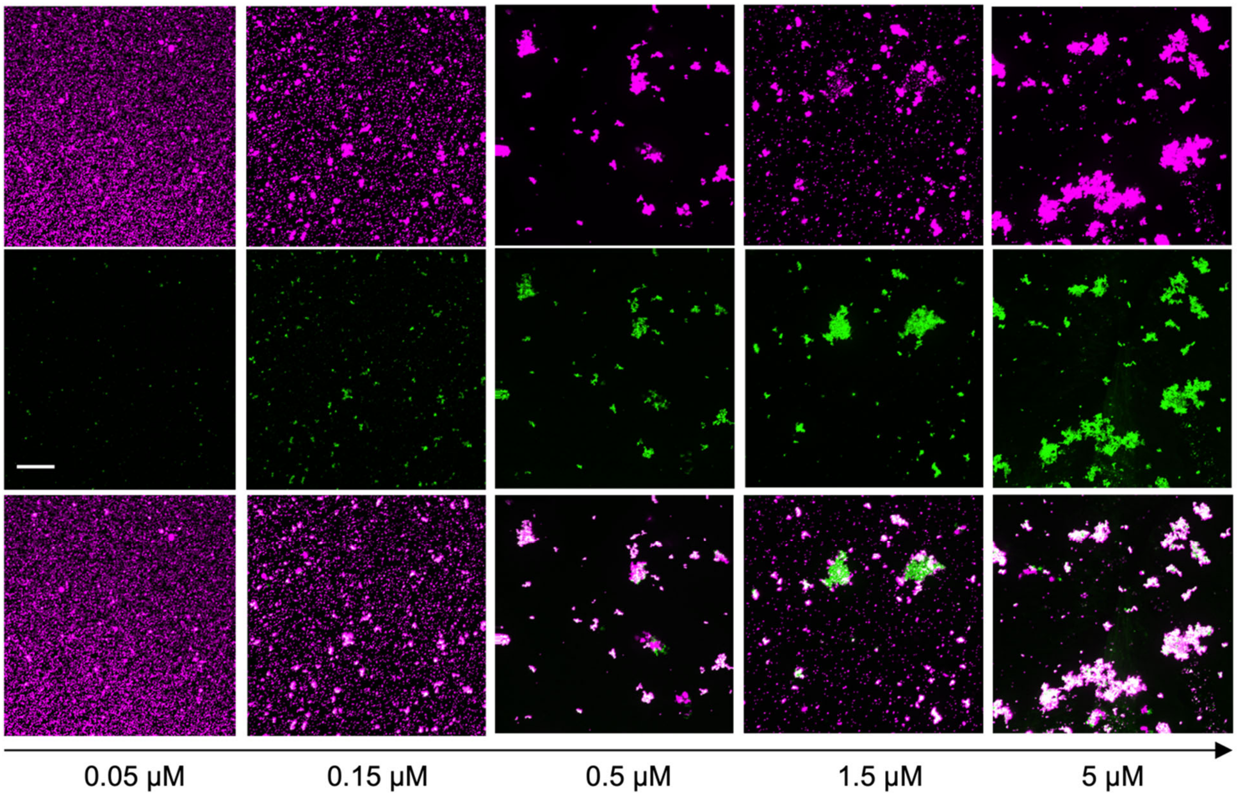


**Fig. S7**: **Maximum projection images of co-aggregates formed at increasing concentrations of FITC-labeled Bap1-57aa.** Shown from top to bottom are the bead channel, the peptide channel, and the overlaid channel, respectively. Scale bar: 20 μm.

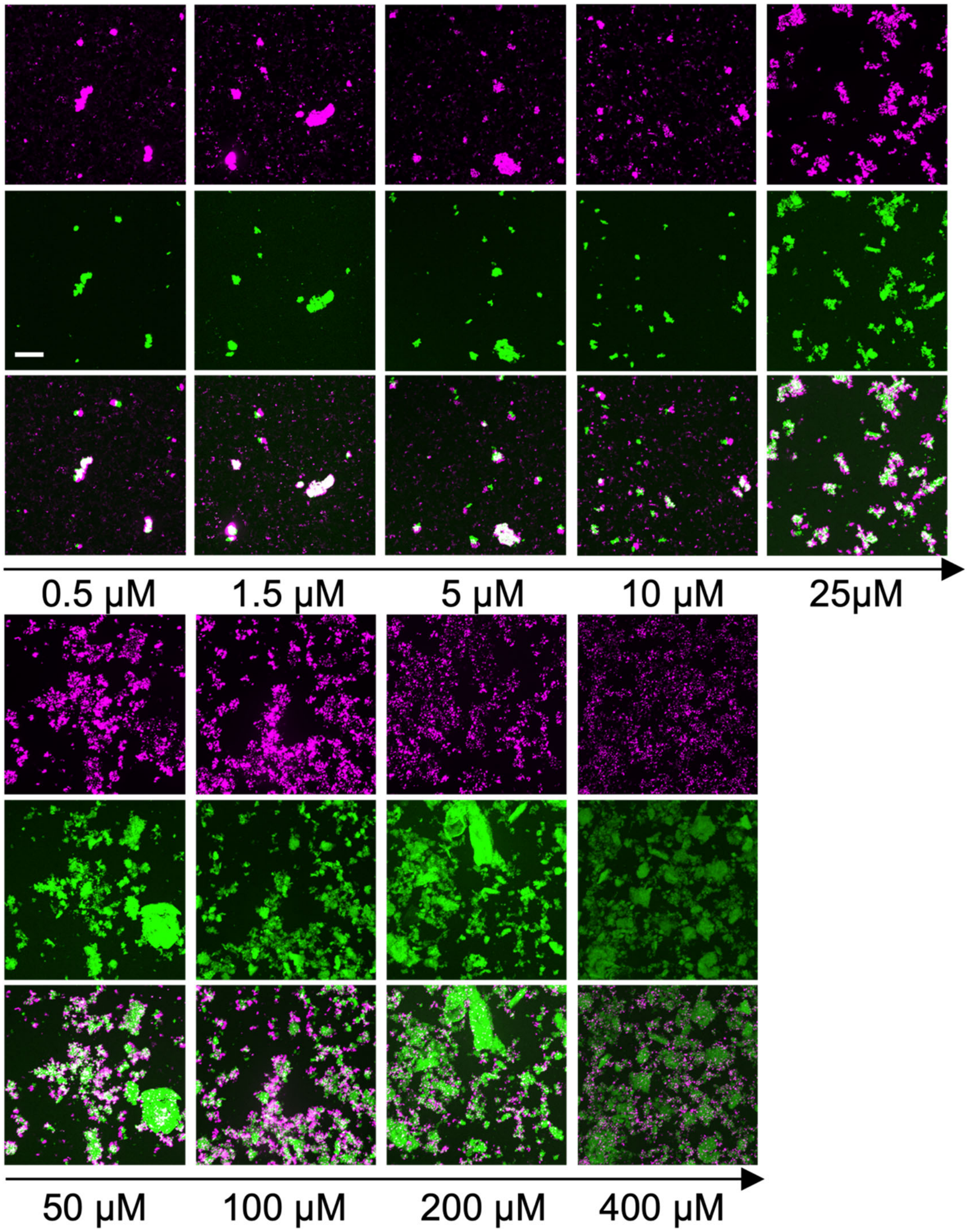


**Fig. S8**: **Maximum projection images of co-aggregates formed at increasing concentrations of *E. coli*-purified Bap1-57aa (labeled with NanoOrange).** Shown from top to bottom are the bead channel, the peptide channel, and the overlaid channel, respectively. Scale bar: 20 µm.

**Table S1: Fitting parameters (±1 standard deviation) for the SFG spectra of the Bap1-57aa peptide deposited on glass substrates using pure water as solvent according to equation 1 (see Materials and Methods).**

| | Parameters | Values | Assignments |
|---|---|---|---|
| Amide I region (Figure 4d, blue) | $\chi_{NR}^{(2)}$ | -0.1 ± 0.003 | N/A |
| | $\omega_1$ ($cm^{-1}$) | 1649.9 ± 0.4 | α-helix |
| | $A_1$ (a.u.) | 3.3 ± 0.1 | |
| | $\Gamma_1$ ($cm^{-1}$) | 20.8 ± 0.6 | |
| | $\omega_2$ ($cm^{-1}$) | 1628.3 ± 0.5 | β-sheet |
| | $A_2$ (a.u.) | 0.5 ± 0.1 | |
| | $\Gamma_2$ ($cm^{-1}$) | 9.8 ± 0.4 | |
| NH/OH stretch region (Figure 4e, blue) | $\chi_{NR}^{(2)}$ | -0.01 ± 0.006 | N/A |
| | $\omega_1$ ($cm^{-1}$) | 3528.8 ± 5.1 | water (OH stretch) |
| | $A_1$ (a.u.) | 31.3 ± 0.9 | |
| | $\Gamma_1$ ($cm^{-1}$) | 145.0 ± 4.2 | |
| | $\omega_2$ ($cm^{-1}$) | 3328.0 ± 1.9 | α-helix (NH stretch) |
| | $A_2$ (a.u.) | 0.93 ± 0.16 | |
| | $\Gamma_2$ ($cm^{-1}$) | 23.9 ± 3.4 | |
| | $\omega_3$ ($cm^{-1}$) | 3246.5 ± 4.0 | β-sheet (NH stretch) |
| | $A_3$ (a.u.) | 0.88 ± 0.15 | |
| | $\Gamma_3$ ($cm^{-1}$) | 35.2 ± 2.8 | |

**Table S2**: **Fitting parameters (±1 standard deviation) for the SFG spectra of the Bap1-57aa peptide deposited on glass substrates using 40% DMSO as solvent according to equation 1 (see Materials and Methods).**

| | Parameters | Values | Assignments |
|---|---|---|---|
| Amide I region (Figure 4d, red) | $\chi_{NR}^{(2)}$ | -0.04 ± 0.003 | N/A |
| | $\omega_1$ (a.u.) | 1682.2 ± 2.0 | β-sheet |
| | $A_1$ (a.u.) | 14.0 ± 0.3 | |
| | $\Gamma_1$ (cm$^{-1}$) | 36.5 ± 1.3 | |
| | $\omega_2$ (cm$^{-1}$) | 1620.8 ± 1.2 | β-sheet |
| | $A_2$ (a.u.) | 2.0 ± 0.4 | |
| | $\Gamma_2$ (cm$^{-1}$) | 19.6 ± 1.9 | |
| NH/OH stretch region (Figure 4e, red) | $\chi_{NR}^{(2)}$ | -0.1 ± 0.008 | N/A |
| | $\omega_1$ (cm$^{-1}$) | 3386.1 ± 9.7 | water (OH stretch) |
| | $A_1$ (a.u.) | 13.1 ± 3.2 | |
| | $\Gamma_1$ (cm$^{-1}$) | 162.0 ± 22.1 | |
| | $\omega_2$ (cm$^{-1}$) | 3269.8 ± 2.9 | β-sheet (NH stretch) |
| | $A_2$ (a.u.) | 2.4 ± 0.4 | |
| | $\Gamma_2$ (cm$^{-1}$) | 50.0 ± 4.3 | |

### Table S3: Bacterial strains used in this study.

| Strain Name in Manuscript | Genotype and Antibiotic Resistance | Description | Strain# & Reference |
|---|---|---|---|
| $\Delta$*rbmC* *bap1*$_{\Delta\beta\text{-prism}}$ | *vpvC*$^{W240R}$, $\Delta$*rbmC*, *bap1*$_{\Delta\beta\text{-prism}}$, $\Delta$*VC1807*::*P*$_{tac}$-*mNeonGreen*, Spec$^{R}$ | Replacement of *bap1*$^{WT}$ with a *bap1* construct lacking the prism but with remaining 57aa by cotransformation with $\Delta$*VC1807*::*P*$_{tac}$-*mNeonGreen,* Spec$^{R}$ into a rugose strain lacking *rbmC* (JY071) | ZJ087 (*34*) |
| $\Delta$*rbmC* *bap1*$_{\Delta\beta\text{-prism}\Delta 57aa}$ | *vpvC*$^{W240R}$, $\Delta$*rbmC*, *bap1*$_{\Delta\beta\text{-prism}\Delta 57aa}$, $\Delta$*VC1807*::*P*$_{tac}$-*mNeonGreen*, Spec$^{R}$ | Replacement of *bap1*$^{WT}$ with a *bap1* construct lacking the β-prism domain and 57aa by cotransformation with $\Delta$*VC1807*::*P*$_{tac}$-*mNeonGreen,* Spec$^{R}$ into a rugose strain lacking *rbmC* (JY071) | ZJ074 (*34*) |
| $\Delta$*rbmC* *bap1*$_{\Delta\beta\text{-prism+57aa*(2-repeat)}}$ | *vpvC*$^{W240R}$, $\Delta$*rbmC*, *bap1*$_{\Delta\beta\text{-prism+57aa*(2-repeat)}}$, $\Delta$*VC1807*::*P*$_{tac}$-*mNeonGreen*, Spec$^{R}$ | Replacement of *bap1*$^{WT}$ with a *bap1* construct lacking the prism with remaining 57aa variant (2-repeat) by cotransformation with $\Delta$*VC1807*::*P*$_{tac}$-*mNeonGreen,* Spec$^{R}$ into a rugose strain lacking *rbmC* (JY071) | XH122 (*44*) |
| $\Delta$*rbmC* *bap1*$_{\Delta\beta\text{-prism+57aa*(core motif)}}$ | *vpvC*$^{W240R}$, $\Delta$*rbmC*, *bap1*$_{\Delta\beta\text{-prism+57aa*(core motif)}}$, $\Delta$*VC1807*::*P*$_{tac}$-*mNeonGreen*, Spec$^{R}$ | Replacement of *bap1*$^{WT}$ with a *bap1* construct lacking the prism with remaining 57aa variant (core motif) by cotransformation with $\Delta$*VC1807*::*P*$_{tac}$-*mNeonGreen,* Spec$^{R}$ into a rugose strain lacking *rbmC* (JY071) | XH166 (*44*) |
| $\Delta$*rbmC* *bap1*$_{\Delta\beta\text{-prism+57aa*(WFFG->LGPE)}}$ | *vpvC*$^{W240R}$, $\Delta$*rbmC*, *bap1*$_{\Delta\beta\text{-prism+57aa*(WFFG->LGPE)}}$, $\Delta$*VC1807*::*P*$_{tac}$-*mNeonGreen*, Spec$^{R}$ | Replacement of *bap1*$^{WT}$ with a *bap1* construct lacking the prism with remaining 57aa variant (WFFG->LGPE) by cotransformation with $\Delta$*VC1807*::*P*$_{tac}$-*mNeonGreen,* Spec$^{R}$ into a rugose strain lacking *rbmC* (JY071) | XH146 (*44*) |
| $\Delta$*rbmC* *bap1*$_{\Delta\beta\text{-prism+C-terminal 57aa}}$ | *vpvC*$^{W240R}$, $\Delta$*rbmC*, *bap1*$_{\Delta\beta\text{-prism+C-terminal 57aa}}$, $\Delta$*VC1807*::*P*$_{tac}$-*mNeonGreen*, Spec$^{R}$ | Replacement of *bap1*$^{WT}$ with a *bap1* construct lacking the prism but containing 57aa at C-terminus by cotransformation with $\Delta$*VC1807*::*P*$_{tac}$-*mNeonGreen,* Spec$^{R}$ into a rugose strain lacking *rbmC* (JY071) | XH169 (*44*) |
| *bap1*$_{\Delta\beta\text{-prism}}$-*3XFLAG* | *vpvC*$^{W240R}$, *bap1*$_{\Delta\beta\text{-prism}}$-*3XFLAG*, $\Delta$*VC1807*::*P*$_{tac}$-*mNeonGreen,* Spec$^{R}$ | Replacement of *bap1*$^{WT}$ with a *bap1* construct lacking the prism but with remaining 57aa and containing a C-terminal 3XFLAG tag by cotransformation with $\Delta$*VC1807*::*P*$_{tac}$-*mNeonGreen,* Spec$^{R}$ into a rugose strain (JY028) | XH024 (*34*) |
| *bap1*$_{\Delta\beta\text{-prism}\Delta 57aa}$-*3XFLAG* | *vpvC*$^{W240R}$, *bap1 bap1*$_{\Delta\beta\text{-prism}\Delta 57aa}$-*3XFLAG*, $\Delta$*VC1807*::*P*$_{tac}$-*mNeonGreen,* Spec$^{R}$ | Replacement of *bap1*$^{WT}$ with a *bap1* construct lacking the β-prism domain and 57aa and containing a C-terminal 3XFLAG tag by cotransformation with | ZJ091 (*34*) |

| | | | |
|---|---|---|---|
| | | $\Delta$*VC1807*::$P_{tac}$-*mNeonGreen,* $Spec^R$ into a rugose strain (JY028) | |
| *bap1*$_{\Delta\beta\text{-}prism+57aa*(core\ motif)}$-*3XFLAG* | *vpvC*$^{W240R}$, *bap1*$_{\Delta\beta\text{-}prism+57aa*(core\ motif)}$-*3XFLAG,* $\Delta$*VC1807*::$P_{tac}$-*mNeonGreen*, $Spec^R$ | Replacement of *bap1*$^{WT}$ with a *bap1* construct lacking the prism with remaining 57aa variant (core motif) and containing a C-terminal 3XFLAG tag by cotransformation with $\Delta$*VC1807*::$P_{tac}$-*mNeonGreen,* $Spec^R$ into a rugose strain (JY028) | XH174 (*44*) |
| *bap1*$_{\Delta\beta\text{-}prism+57aa*(WFFG\text{-}>LGPE)}$-*3XFLAG* | *vpvC*$^{W240R}$*, bap1*$_{\Delta\beta\text{-}prism+57aa*(WFFG\text{-}>LGPE)}$-*3XFLAG,* $\Delta$*VC1807*::$P_{tac}$-*mNeonGreen*, $Spec^R$ | Replacement of *bap1*$^{WT}$ with a *bap1* construct lacking the prism with remaining 57aa variant (WFFG-LGPE) and containing a C-terminal 3XFLAG tag by cotransformation with $\Delta$*VC1807*::$P_{tac}$-*mNeonGreen,* $Spec^R$ into a rugose strain (JY028) | XH144 (*44*) |

83
